# Chemical proteomics identifies signal peptidase IB (SpsB) as a target of the SOS response inhibitor OXF-077 and a regulator of quinolone resistance emergence in *Staphylococcus aureus*

**DOI:** 10.1101/2025.11.21.689737

**Authors:** Jacob D. Bradbury, Timothy R. Walsh, Thomas Lanyon-Hogg

**Affiliations:** Department of Pharmacology, University of Oxford, OX1 3QT, UK; Ineos Oxford Institute for Antimicrobial Research, University of Oxford, OX1 3RE, UK

**Keywords:** Antibiotics, Photoaffinity labelling, SOS response, Chemical proteomics, Antimicrobial resistance

## Abstract

Antimicrobial resistance (AMR) is an existential threat to health globally, and novel compounds that act through new targets are urgently required to combat increasing levels of resistance. One emerging strategy is the development of ‘antibiotic adjuvants’ that can slow the evolution of resistance by inhibiting the mutagenic SOS response in bacteria, in order to prolong the clinical lifetime of antibiotics. **OXF-077** is a potent SOS response inhibitor that suppresses the emergence of ciprofloxacin resistance in *Staphylococcus aureus*; however, the cellular target of **OXF-077** is unknown. We report here the use of affinity-based protein profiling to identify signal peptidase IB (SpsB) as a target of **OXF-077**. Genetic and chemical studies demonstrated that SpsB is required for upregulation of the SOS response gene *recA*, increased frequency of resistance emergence to ciprofloxacin, and activation of the mutagenic SOS response in *S. aureus*. SpsB is therefore postulated to regulate the quinolone-activated SOS response in *S. aureus*, and can be targeted by small-molecule inhibitors such as **OXF-077** to slow the evolution of resistance. Collectively, this work delivers SpsB as an attractive new drug target for the development of antibiotic adjuvants to combat the urgent threat of AMR.

**Graphical abstract:** 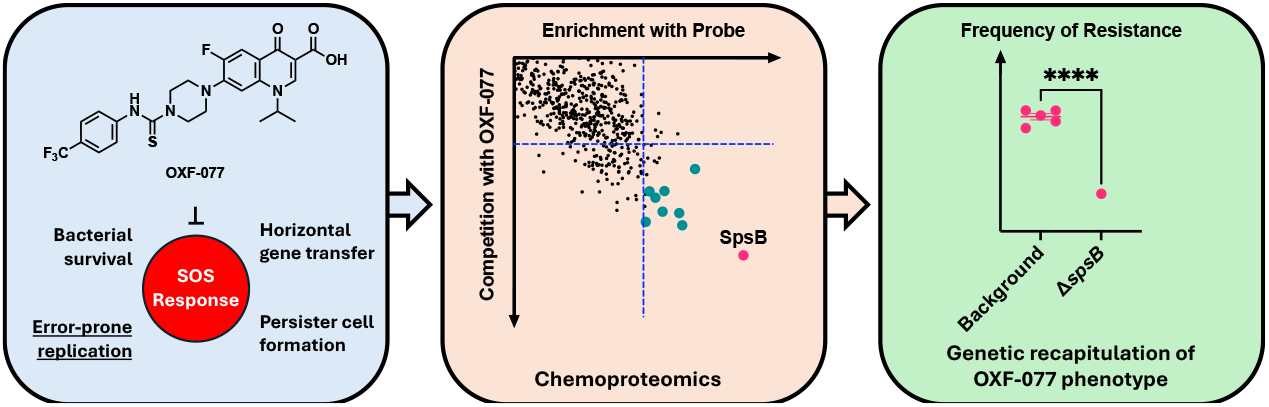

## Introduction

Antimicrobial resistance (AMR) is one of the greatest challenges to health globally. AMR was associated with an estimated 4.95 million deaths in 2019, of which 1.27 million deaths were directly attributable to bacterial AMR.^[1]^ Antibiotics are essential for the life expectancy made possible by advancements in modern medicine, being required for surgery, cancer chemotherapy, childbirth, and the treatment of minor infections that could otherwise be life-threatening.^[2]^ AMR is accelerated by the use and overuse of existing antibiotics in both humans and agriculture,^[3]^ and the efficacy of new antibiotics is rapidly compromised by the rapid emergence of novel resistance mechanisms or the selection of pre-existing resistance mechanisms.^[4]^ This rapid emergence of AMR means new antibiotics must be protected to prolong their clinical effectiveness; however, this safeguarding also limits the commercial market for any new antibiotic product and therefore disincentivises development.^[5]^

One aspect of addressing the challenge of AMR is the development of new compounds with novel mechanisms of action. This includes both direct-acting antibiotics from new chemical classes, as well as ‘antibiotic adjuvants’ that can be paired with antibiotics to block resistance mechanisms and resensitize bacteria, such as ß-lactamase inhibitors.^[6]^ An emerging strategy is the development of compounds to reduce the rate of resistance evolution, for example by inhibiting the SOS response in bacteria (Fig. 1A).^[7,8]^ The SOS response is activated by bacterial DNA-damage, which is caused by treatment with fluoroquinolone antibiotics such as ciprofloxacin (**CFX, 1**, Fig. 1B). DNA double-strand breaks from **CFX** treatment are processed by the helicase–nuclease enzymes AddAB or RecBCD to generate a 3’ single-stranded DNA (ssDNA) overhang which is recognised by the protein RecA to form a RecA-ssDNA nucleoprotein filament (RecA*).^[9,10]^ RecA* initiates homologous recombination and DNA repair, and also interacts with the transcriptional repressor protein LexA causing auto-cleavage of LexA and expression of SOS box genes.^[9]^ The SOS box genes include error-prone polymerases that increase the rate of mutation, as well as genes promoting horizontal gene transfer and persister cell formation (Fig 1A).^[11]^ The range of resistance mechanisms activated by the SOS response makes this pathway an attractive target for the development of antibiotic adjuvants to slow resistance emergence; however, to-date no SOS inhibitors have progressed to the clinic.^[11]^

**Figure 1.**
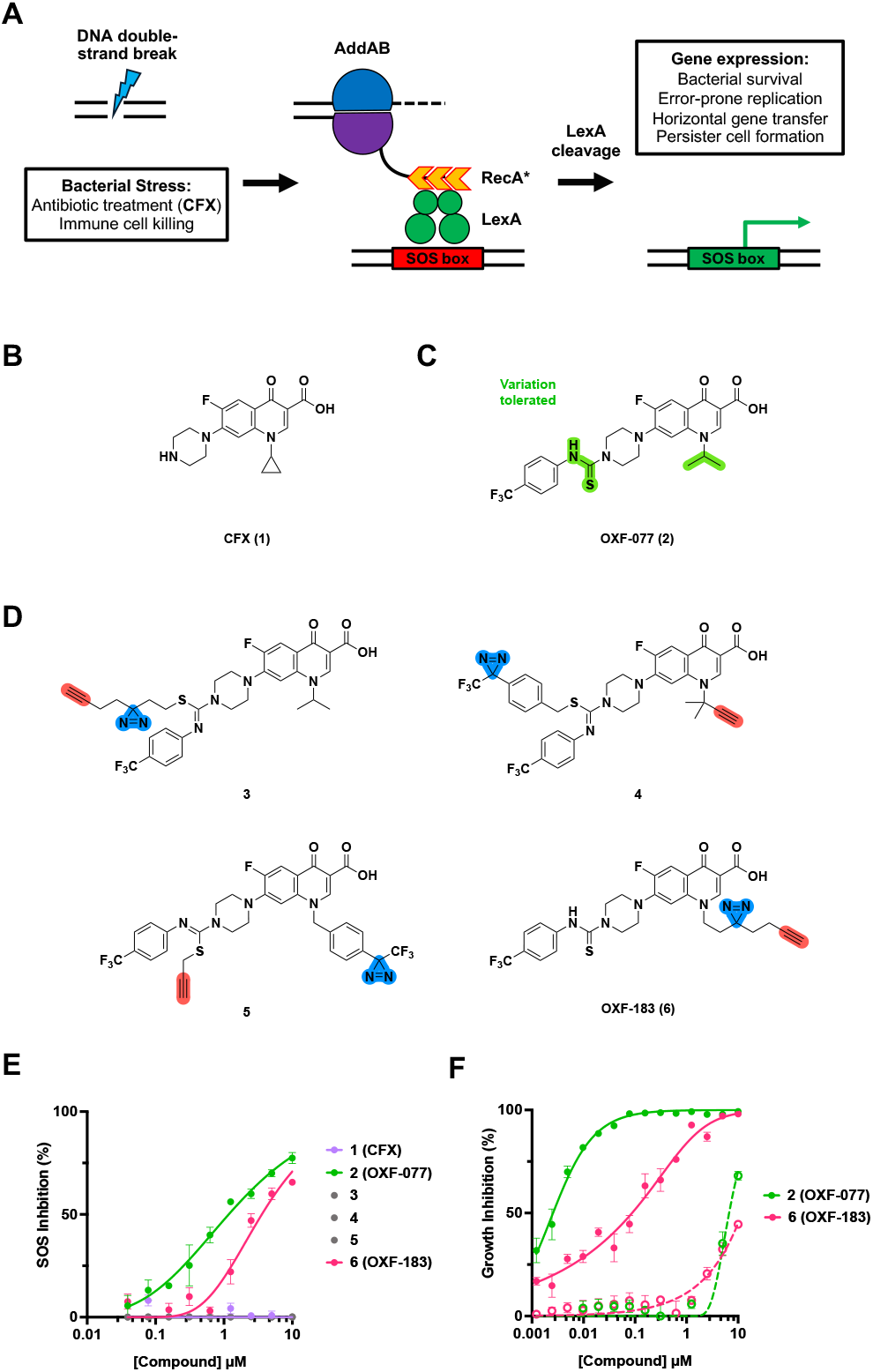
Development and validation of photochemical probes of OXF-077, an inhibitor of the *S. aureus* SOS response pathway. **(A)** Schematic of SOS response pathway and associated gene expression.^[11]^ **(B)** Structure of DNA-damaging quinolone antibiotic, ciprofloxacin (**CFX, 1**). **(C)** Structure of SOS response inhibitor, **OXF-077** (**2**), with structural positions that tolerate variation highlighted.^[12]^ **(D)** Structures of probes **3**-**6** used in this study. **(E)** SOS response inhibition (%) of **CFX, OXF-077** and probes **3**-**6** in MRSA JE2 SOS reporter assay. **(F)** Growth inhibition (%) of **OXF-077** and **OXF-183** (**6**) in the presence of (solid line) or absence of (dashed line) of **CFX** (6 µM). Data represent mean ± standard error of the mean (SEM), n=3 independent biological replicates.

**OXF-077** (**2**, Fig. 1C) is an inhibitor of the DNA repair and SOS response pathways in methicillin-resistant *Staphylococcus aureus* (MRSA).^[12]^ **OXF-077** was developed through a phenotypic structure-activity relationship (SAR) study of putative AddAB/RecBCD inhibitors **ML-328** and **IMP-1700** (Fig. S1).^[7,8,12]^ **OXF-077** reduces the rate of **CFX** resistance emergence in *S. aureus* and also re-sensitizes *S. aureus* to concentrations of **CFX** below the clinical breakpoint once resistance has emerged.^[12]^ However, despite the therapeutically-beneficial phenotypes produced by **OXF-077** in slowing or reversing **CFX** resistance, the cellular target(s) of this series remains the subject of debate.^[12,13]^ Identification of the target of **OXF-077** in MRSA is critical step to guide further translation development of this promising series of SOS inhibitors.

We report here the use of affinity-based protein profiling (A*f*BPP) to identify the cellular targets of **OXF-077** in live MRSA. A*f*BPP identified multiple **OXF-077** target proteins, including signal peptidase IB (SpsB) which was shown for the first time to regulate expression of the SOS response gene *recA* and *S. aureus* frequency of resistance to **CFX**. This work therefore delivers SpsB as a novel and ligandable target to inhibit the SOS response in *S. aureus* and slow the evolution of AMR.

## Results

### Development of an OXF-077 affinity-based probe

In order to identify the cellular targets of **OXF-077**, a panel of four photochemical probes were designed containing a diazirine for covalent protein crosslinking upon UV irradiation, and an alkyne handle for biorthogonal attachment of reporter groups using copper-catalysed alkyne−azide cycloaddition (CuAAC) ‘click chemistry’ (Fig. 1D). Probes incorporated the diazirine and alkyne groups at **OXF-077** sites where prior SAR data indicates variation can be tolerated whilst maintaining SOS inhibition, specifically the isopropyl and thiourea groups.^[12]^ Probe **3** was prepared by direct alkylation of **OXF-077** with a minimalist photocrosslinkable-clickable handle (Scheme S1A).^[14]^ Synthesis of probes **4** - **6** built on intermediates synthesised as A*f*BPP probes of **CFX**.^[15]^ Probe **4** was synthesised from a 3-methylbut-1-yne **CFX** analogue coupled with 4-trifluoromethyl isothiocyanate, and subsequently alkylated at the thiourea with 3-(4-(bromomethyl)phenyl)-3-(trifluoromethyl)-diazirine (Scheme S1B). Probe **5** was similarly synthesised from a 3-(4-(methyl)phenyl)-3-(trifluoromethyl)-diazirine **CFX** analogue, 4-trifluoromethyl isothiocyanate and propargyl bromide (Scheme S1B). Finally, probe **6** was prepared from a **CFX** analogue containing a minimalist photocrosslinkable-clickable handle^[14]^ at the isopropyl position coupled with 4-trifluoromethyl isothiocyanate (Scheme S1B).

To assess the ability of probes to recapitulate the biological activity of **OXF-077**, SOS response inhibition was assessed in a JE2 MRSA cellular reporter assay expressing GFP under control of the *recA* promoter, *precA-gfp*.^[16]^ **OXF-077** inhibited the **CFX**-activated SOS response with an IC_50_ of 1.2 ± 0.1 µM (Fig. 1E), consistent with previous reports.^[12]^ Only compound **4** exhibited SOS response inhibition, with an IC_50_ of 3.6 ± 0.2 µM (Fig. 1E). SOS activation by probe **6** or **OXF-077** was also assessed, with neither compound activating the SOS response <10 µM (Fig. S2A).

In addition to inhibiting the SOS response, **OXF-077** potentiates the activity of **CFX** below the minimum inhibitory concentration (MIC) in JE2 MRSA (**CFX** MIC = 8 µg mL^-1^, 24 µM).^[12]^ Growth inhibition by **OXF-077** and probe **6** was measured as single agents or in combination with a sublethal dose of **CFX** (6 μM). **OXF-077** potentiated the activity of **CFX**, exhibiting growth inhibition IC_50_ with **CFX** co-treatment of 3.0 ±0.7 nM, compared to an independent growth inhibition IC_50_ as a single agent of 6.7 ± 0.2 µM (Fig. 1F). Similarly, probe **6** also potentiated the activity of **CFX**, displaying an independent IC_50_ >10 μM and a combination with **CFX** IC_50_ of 88 ± 5 nM (Fig. 1F). Having recapitulated the biological effects of **OXF-077** in SOS inhibition and DNA-damage potentiation, probe **6**, which we here term **OXF-183** was progressed to target profiling studies.

### Identification of OXF-077 protein targets via affinity-based protein profiling

Having validated **OXF-183** as a suitable probe for identifying targets of **OXF-077**, photoactivation of **OXF-183** upon UV irradiation (365 nm) was assessed over a 4-min time course and quantified by HPLC. The half-life of **OXF-183** was 17 s (95% CI = 13 to 22 s) indicating rapid activation of the diazirene group by UV light (Fig. S2B); in contrast, **OXF-077** was unaffected after 4 min of UV irradiation.

Next, protein labelling in live JE2 MRSA was investigated using an in-gel fluorescence readout. Overnight cultures were washed and treated with DMSO, **OXF-183**, or **OXF-183** in competition with **OXF-077** (Fig. 2A). Samples were then irradiated for 1 min to induce photocrosslinking, lysed, and the lysate protein concentration normalised. The probe-modified proteins were functionalised by a CuAAC ‘click’ reaction with the trifunctional capture reagent Azido-TAMRA-biotin (AzTB). AzTB contains a TAMRA fluorophore for in-gel visualisation of labelled proteins and a biotin affinity tag for enrichment of labelled proteins with neutravidin agarose resin (Fig. S3A). Probe-labelled proteins were separated by SDS-PAGE and analysed by in-gel fluorescence scanning, with total protein staining used to assess loading. Protein labelling was both probe- and UV-dependent (Fig. S3B), with labelling increasing in a dose-dependent manner (Fig. S3C). Co-treatment of cells with **OXF-077** (1, 10, and 100 µM) and **OXF-183** (2 µM) resulted in competition for specific labelled proteins (Fig. 2B), such as at ∼120 kDa (band A) and ∼25 kDa (band B), suggesting these proteins may be targets of **OXF-077**.

**Figure 2.**
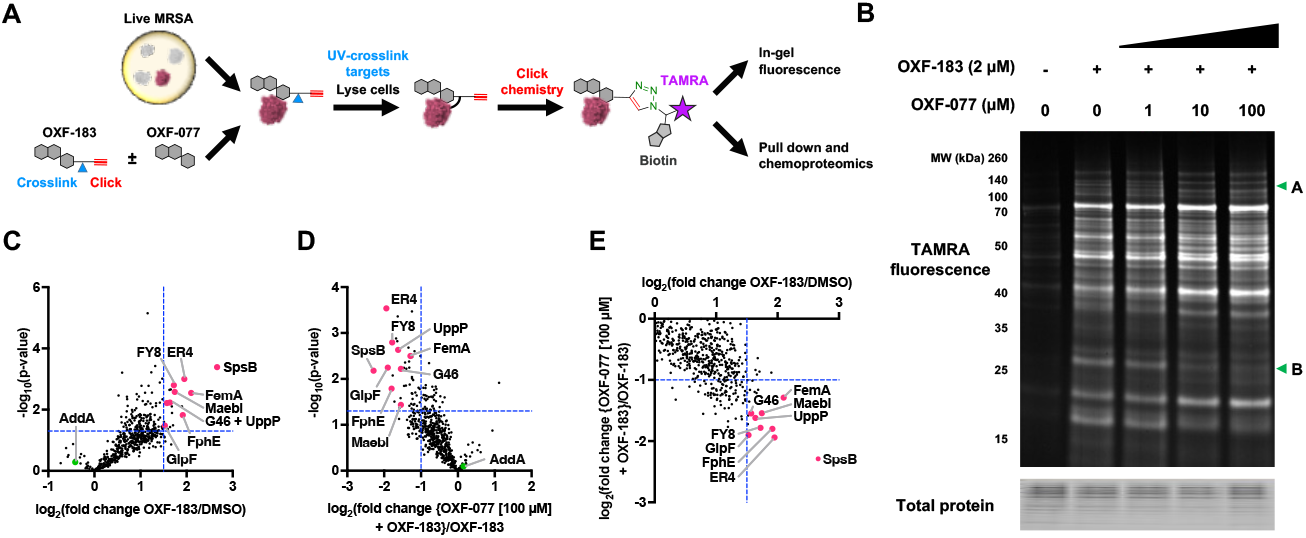
Affinity-based protein profiling (A*f*BPP) with OXF-077 photocrosslinking probe OXF-183. **(A)** A*f*BPP workflow for target protein labelling by **OXF-183** and subsequent functionalisation for analysis. **(B)** In-gel fluorescence of protein labelling in live MRSA using **OXF-183** in competition with **OXF-077**. Two prominent protein bands (A and B) which displayed competition with **OXF-077** are annotated. **(C)** Proteomic analysis of proteins enriched following **OXF-183** (2 µM) treatment compared to DMSO. **(D)** Proteomic analysis of proteins enriched following **OXF-183** (2 µM) treatment in the presence or absence of **OXF-077** (100 µM) competition. **(E)** Comparison of fold-change of proteins enriched by **OXF-183** (2 µM) treatment with fold-change of proteins competed by **OXF-077** (100 µM) co-treatment. Data represent mean ± standard error of the mean (SEM), n=3 independent biological replicates.

To identify and quantify the binding partners of **OXF-077** on a whole-proteome scale, the protein targets were analysed by chemical proteomics. Overnight cultures were treated with DMSO, **OXF-183** (2 µM) alone, or **OXF-183** (2 µM) in competition with **OXF-077** (10 µM or 100 µM), UV crosslinked, and functionalised with AzTB. Labelled proteins were enriched by pull-down with NeutrAvidin agarose, washed, digested on-bead with LysC and trypsin, and the resulting peptides analysed by tandem LC-MS/MS.

29 proteins showed statistically significant enrichment (*p*-value <0.05, log_2_(fold change) >1.5) following treatment with **OXF-183** compared to DMSO (Fig. 2C, Table S1). Competition for protein labelling with **OXF-077** (10 and 100 µM) resulted in a statistically significant decrease (*p*-value <0.05, log_2_(fold change) <-1) in 4 proteins at 10 µM (Fig. S3D, Table S1) **OXF-077** and 50 proteins at 100 µM **OXF-077** (Fig. 2D, Table S1). Nine proteins showed both statistically significant enrichment by **OXF-183** and competition by **OXF-077** at 100 µM (Fig. 2E, Table 1, Table S1) and were therefore selected for further investigation as potential targets of **OXF-077**.

**Table 1.**
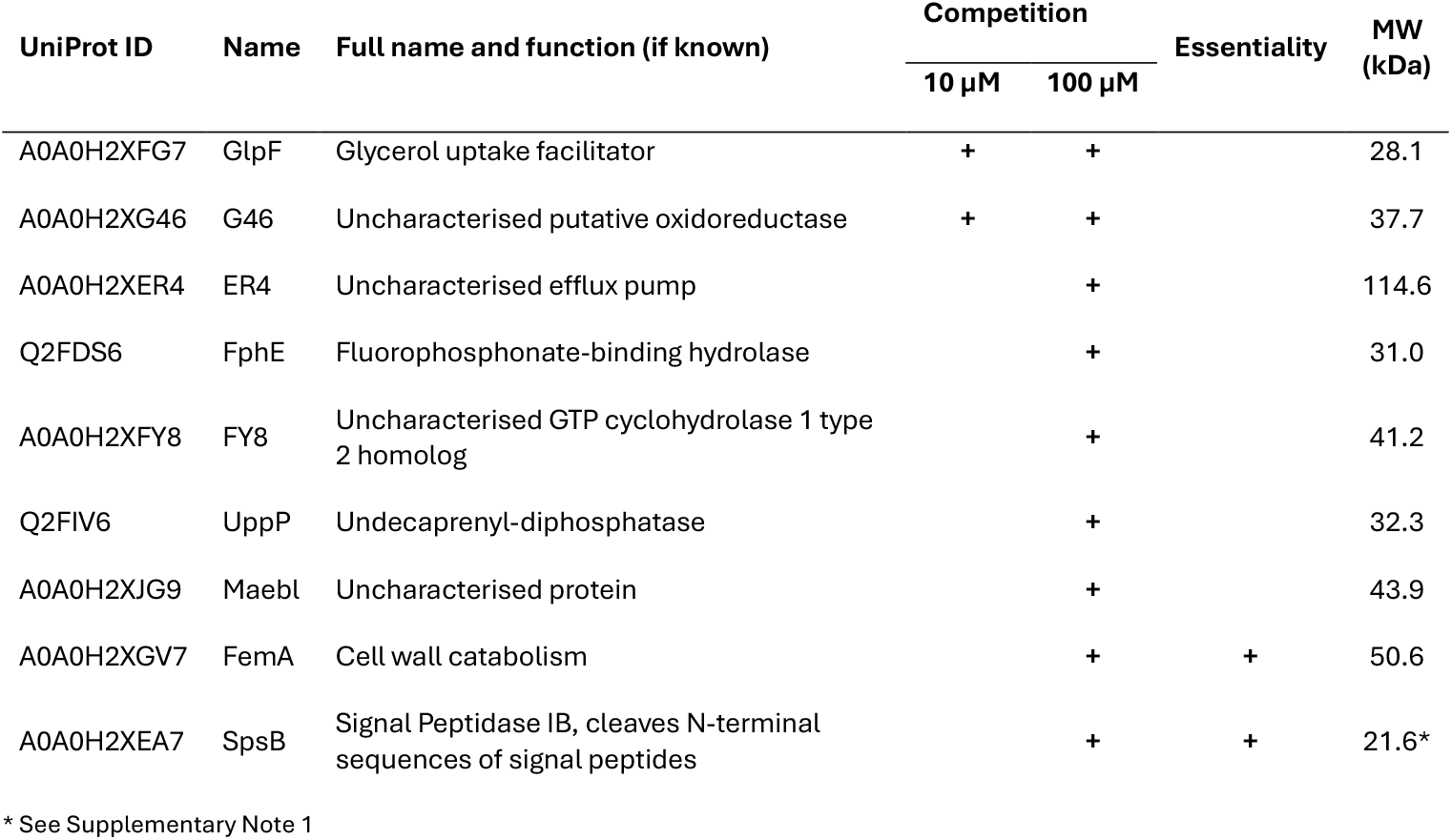
Proteins identified by chemoproteomics.

Of the functionally annotated proteins in the **OXF-077** target profile, GlpF showed competition at 10 µM and 100 µM, whereas FphE, UppP, FemA and SpsB showed competition at 100 µM only (Table 1). Signal Peptidase IB (SpsB) had the highest **OXF-183** enrichment and highest **OXF-077** (100 µM) competition of any protein (Fig. 2E). Interestingly, the AddA subunit of the proposed **OXF-077** target AddAB was detected in these experiments; however, AddA did not reach the threshold for statistically significant enrichment or competition by **OXF-077** (Fig. 2C,D).

### Validation of putative OXF-077 targets

Having identified a list of putative targets for **OXF-077** by chemoproteomics, knockout mutants were used to validate engagement of these proteins by **OXF-183**. Transposon-insertion mutants are available for the seven non-essential protein targets in the JE2 strain.^[17]^ Additionally, an allelic replacement knockout mutant of *spsB* is available in the RN1 *S. aureus* strain,^[18]^ where knockout of the transcriptional repressor *cro/cI* circumvents the essentiality of *spsB* by overexpression of a putative ABC transporter.^[19]^ Only one potential target, the essential protein FemA, did not have any corresponding knockout strains and as **OXF-077** does not inhibit cell viability when dosed as a single agent, FemA was not investigated further.

Initially, knockout mutants of the potential targets in JE2 MRSA were investigated by A*f*BPP with **OXF-183** and in-gel visualisation using Azido-TAMRA (AzT, Fig. S4A). AfBPP in the mutant strains validated two proteins, FY8 and ER4, as binding partners of **OXF-183**, with ER4 corresponding to band A at ∼120 kD and FY8 corresponding to a band at ∼45 kDa not competed by **OXF-077** (Fig. S4B). ER4 is annotated as an uncharacterised efflux pump, and therefore represented a potential target that could mediate the potentiation of cell-killing by **CFX**. However, no difference in **CFX** tolerance or **CFX** potentiation by **OXF-077** (5 µM) was observed with ER4 knockout, or in any other available MRSA JE2 target knockout strains (Fig. S4C,D). SOS activation by **CFX** induces the formation of persister cells,^[11]^ and treatment of MRSA JE2 with **OXF-077** (5 µM) resulted in a decrease in the number of persister cells (Fig. S4E); however, again no reduction in persister cell formation was observed for any knockout strain (Fig. S4F). Collectively these data indicate that whilst FY8 and ER4 are engaged by **OXF-183**, these proteins are not responsible for the phenotypic effects of **OXF-077**.

The remaining **OXF-077** binding protein, SpsB, was therefore investigated the in RN1 *S. aureus* strain. In-gel A*f*BPP identified SpsB as the **OXF-077** target band B at ∼25 kDa, which showed reduced fluorescent labelling with *spsB* knockout (Fig. 3A, Fig. S5A). The ability of SpsB to control **CFX** resistance and **OXF-077** potentiation was therefore investigated by assessing the MIC of **CFX** alone and in combination with **OXF-077** (5 µM) in WT, Δ*cro/cI* and, Δ*cro/cI* Δ*spsB* strains. All strains possessed the same MIC when treated with **CFX** alone (0.19 µg mL^-1^, 0.063 µM) or **CFX** in combination with 5 µM **OXF-077** (0.048 µg mL^-1^, 16 nM, Fig. 3B). However, Δ*cro/cI* Δ*spsB* exhibited decreased tolerance to **CFX** at sub-MIC concentrations in combination with **OXF-077** (Fig. 3B), suggesting that a second target of **OXF-077** controls DNA-damage repair.

**Figure 3.**
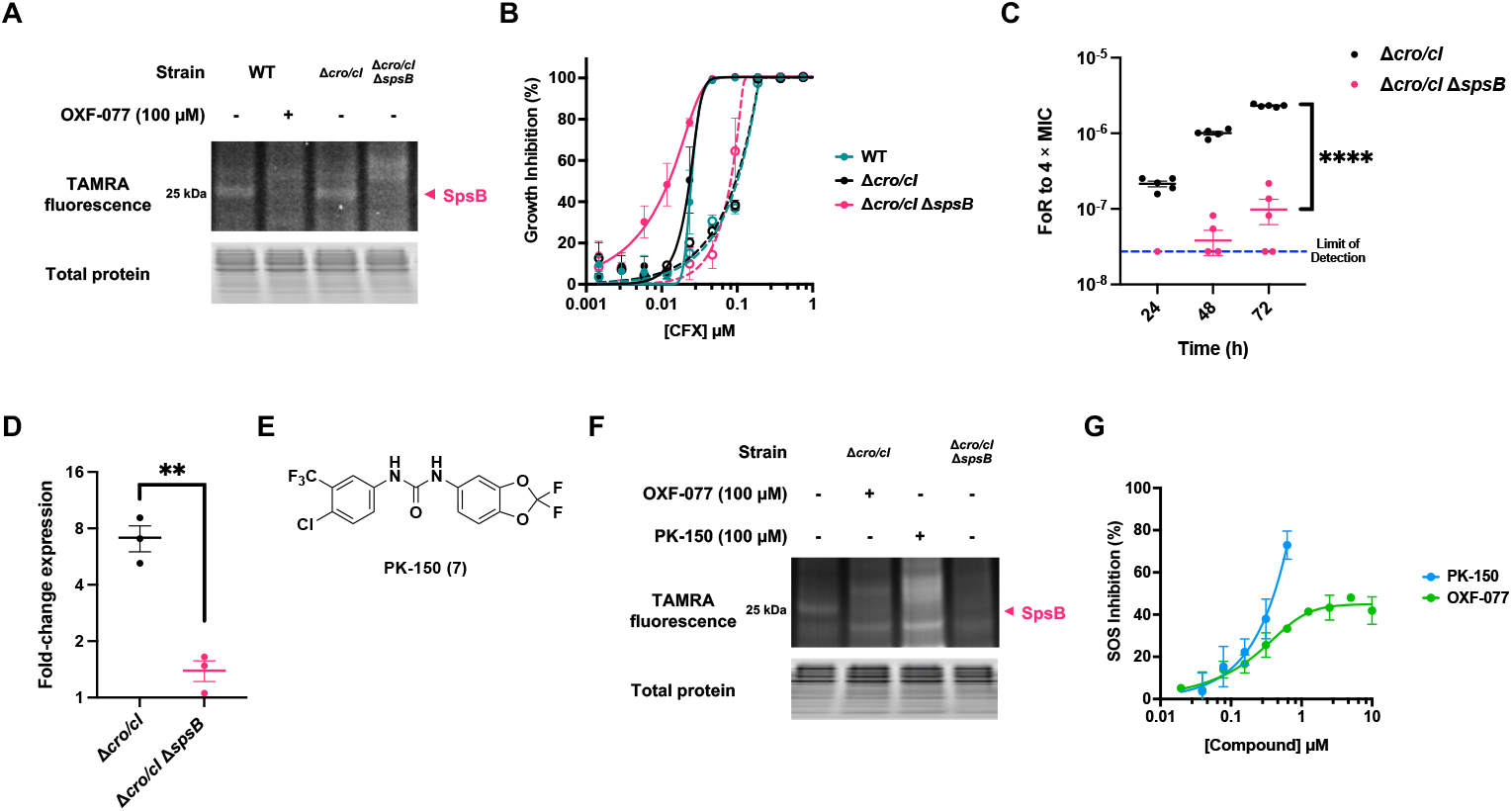
Validation of SpsB as a target of OXF-077 that regulates the SOS response. **(A)** In-gel A*f*BPP **OXF-183** (2 µM) in wild-type (WT) with and without competition of **OXF-077** compared with Δ*cro/cI* and Δ*cro/cI* Δ*spsB*. **(B) CFX** growth inhibition (%) of RN1 strains in the presence (solid line) or absence (dashed line) of **OXF-077** (5 µM). **(C)** Frequency of resistance (FoR) of Δ*cro/cI* and Δ*cro/cI* Δ*spsB* to 4 × MIC of **CFX** over a 72 h time course. Data points not shown are below the limit of detection of one colony. **(D)** Relative fold change in *recA* expression in Δ*cro/cI* and Δ*cro/cI* Δ*spsB* upon treatment with 4 × MIC **CFX. (E) PK-150** (**7**). **(F)** In-gel A*f*BPP with **OXF-183** (2 µM) in Δ*cro/cI* strain with and without competition of **OXF-077** and **PK-150** compared with Δ*cro/cI* Δ*spsB*. The increased lane fluorescence of cells treated with **PK-150** possibly represents an increase in cell permeation of **OXF-183** under these treatment conditions. **(G)** Normalised SOS response inhibition of **PK-150** and **OXF-077** in p*recA-gfp* JE2 *S. aureus*. Data represent mean ± SEM, n≥3 independent biological replicates.

SOS response activation by **CFX** also upregulates mutagenic DNA repair,^[11]^ which can be investigated by measuring resistance evolution in frequency of resistance (FoR) experiments.^[20]^ FoR to **CFX** was therefore assessed in Δ*cro/cI* and Δ*cro/cI* Δ*spsB* strains after 24, 48 and 72 h treatment with **CFX** at 4 × MIC (0.75 µM). The **CFX** FoR was reduced in Δ*cro/cI* Δ*spsB* compared to the Δ*cro/cI* background at all timepoints, and only after 72 h did all 5 biological replicates of Δ*cro/cI* Δ*spsB* possess a FoR above the limit of detection (*p* < 0.00001 at t=72 h, unpaired t-test, Fig. 3C). Activation of the SOS response also triggers expression of the RecA DNA repair protein,^[21]^ therefore *recA* mRNA expression levels were measured by RT-qPCR in Δ*cro/cI* and Δ*cro/cI* Δ*spsB* cells following treatment with **CFX** (4 × MIC) for 2 h. **CFX** treatment caused an increase in *recA* expression in the Δ*cro/cI* background (7.1 ± 1.1-fold change), which was significantly reduced for Δ*cro/cI* Δ*spsB* (1.4 ± 0.18-fold change, *p* = 0.007 unpaired t-test, Fig. 3D). However, *recA* expression was higher in untreated Δ*cro/cI* Δ*spsB* compared to the Δ*cro/cI* background (Fig. S5B), suggesting that Δ*cro/cI* Δ*spsB* may have been under elevated basal stress. Comparison of growth rates of the RN1 strains demonstrated Δ*cro/cI* Δ*spsB* had a longer lag time than Δ*cro/cI* or WT cells (Fig. S5C). Taken together, these data indicated that SpsB knockout recapitulated several of the observed phenotypes produced by **OXF-077** treatment, and suggested that SpsB may be the target of **OXF-077** that modulates the evolution of resistance to **CFX** *via* the SOS response.

SpsB is an extracellularly-bound membrane protein that cleaves the N-terminal of immature proteins following transmembrane secretion.^[22,23]^ Other small-molecule ligands of SpsB have been reported in the literature, such as arylomycin, an SpsB inhibitor,^[24]^ and **PK-150** (**7**, Fig. 3E),^[25]^ an SpsB activator. **PK-150** is a sorafenib analogue that was identified as a SpsB ligand during a kinase inhibitor repurposing campaign.^[25]^ The structure of **PK-150** is similar to key pharmacophoric groups in **OXF-077**, possessing a para-substituted, electron-deficient phenyl (thio)urea motif.^[8,12]^ Binding-site identification via photocrosslinking demonstrates that **PK-150** binds at an allosteric site on SpsB comprised of a hydrophobic cleft close to the substrate peptide-binding groove.^[26]^ **PK-150** was therefore assessed by in-gel A*f*BPP with **OXF-183** which demonstrated that these compounds competed for binding to SpsB (Fig. 3F, Fig. S5D) and suggested that **PK-150** may elicit a similar phenotypic response to **OXF-077** treatment in MRSA. Lastly, therefore, **PK-150** was tested for SOS response inhibition in the JE2 MRSA reporter assay, which demonstrated that **PK-150** was indeed a potent inhibitor of the SOS response in MRSA (Fig. 3G) and also a more potent inhibitor of cell growth than **OXF-077** (Fig. S5E), potentially consistent will the essential nature of SpsB in MRSA. Collectively, these chemical and genetic lines of evidence support a newly identified role for SpsB in regulation of the SOS-mediated antibiotic resistance responses of MRSA.

## Discussion

The emergence of AMR is one of the most serious healthcare challenges of our time, which is already causing >5 million deaths per year and is only predicted to grow.^[1]^ The SOS response controls hypermutation and resistance evolution during antibiotic stress, and is therefore an attractive target for the development of novel adjuvants to combat AMR.^[11]^ **OXF-077** is a potent inhibitor of the SOS response in *S. aureus* that slows and reverses the evolution of resistance to quinolone antibiotics;^[12]^ however, the cellular targets underlying these therapeutically-beneficial **OXF-077** phenotypes were previously unknown. Here, target-agnostic A*f*BPP with photochemical probe **OXF-183** was used to identify signal Peptidase IB (SpsB) as the target of **OXF-077** responsible for SOS inhibition in MRSA. Genetic studies demonstrated that SpsB levels affected the expression of SOS response gene *recA* and the frequency of resistance emergence to **CFX** in *S. aureus* (Fig. 3C,D). SpsB is therefore postulated to act as a novel regulator of the quinolone-activated SOS response in *S. aureus*, with **OXF-077** inhibiting this process. Furthermore, literature SpsB ligand **PK-150**^[25]^ also competed with **OXF-183** for SpsB binding and inhibited the SOS response (Fig. 3F,G), showing that SpsB is not simply a promiscuous target identified in chemoproteomic experiments. As SpsB is an essential protein, this data collectively presents the exciting possibility that SpsB may represent a new antibiotic drug target that can simultaneously elicit both bacterial killing and inhibition of the mutagenic SOS response pathway. Indeed, **PK-150** is a potent inhibitor of cell growth but also shows no detectable resistance development, even in the presence of the mutagen ethyl methanesulfonate.^[25]^ Moreover, SpsB is also an extracellular membrane-bound protein, which therefore circumvents one of the main challenges of cell penetration faced by inhibitors of other intracellular SOS pathway or antibiotic targets.

**OXF-077** potentiated **CFX** growth inhibition even in the Δ*spsB* strain (Fig. 3B), suggesting that a second, SpsB-independent, target for **OXF-077** may mediate the DNA-damage potentiation effect exhibited by this series. Deletion of *spsB* disrupts cell-wall biogenesis and may result in increased **OXF-077** cell penetration,^[27]^ thereby potentially increasing engagement of this secondary intracellular target. A possible candidate for this target identified by A*f*BPP with **OXF-183** is FemA, which is an essential cytoplasmic enzyme involved in peptidoglycan cross-bridge synthesis whose inhibition increases **CFX** susceptibility two- to four-fold.^[28]^ The inhibition of SOS response and DNA-damage potentiation pathways by secondary targets aside from SpsB is also consistent with prior SAR data, which demonstrate that an ethyl ester analogue of **IMP-1700** could strongly inhibit SOS induction despite showing no observable DNA-damage potentiation.^[12]^ SpsB is canonically associated with *S. aureus* quorum sensing^[29]^ and virulence;^[19]^ however, recent evidence suggests that SpsB has a wider role in *S. aureus* physiology, with implications in cell cycle progression and cell envelope biogenesis.^[27]^ SpsB has not previously been implicated in the SOS response; however, SpsB exhibits sequence and functional homology with the SOS repressor protein LexA,^[23]^ and neighbours *addB* in the *S. aureus* genome,^[30]^ supporting a previously unrecognised functional relationship between these proteins in SOS activation. The precise mechanism by which DNA-damage, the SOS response and SpsB function affect the evolution of resistance in MRSA therefore remains a critical question to be answered.^[31]^

Collectively, this work demonstrates that SpsB is a new extracellular and ligandable drug target that can both induce cell killing and disrupt the resistance SOS responses of MRSA. This delivers multiple chemical tools to study SpsB activity and advance therapeutic development of new molecules to combat the global challenge of AMR.

## Acknowledgements

This research was supported by the Wellcome Trust [317713/Z/24/Z and 218514/Z/19/Z] and the Ineos Oxford Institute for Antimicrobial Research. The authors thank Dr. Vaishnavi Ravikumar and Dr. Marjorie Fournier (Advanced Proteomics Facility, Department of Biochemistry, University of Oxford) for assistance with proteomic experiments and analysis, and Prof. Tom W. Muir and Dr. Ben Buchmuller (Department of Chemistry, University of Princeton) for the gift of the RN1 *S. aureus* strains. All authors acknowledge the provision of strains by the Network on Antimicrobial Resistance in *Staphylococcus aureus* (NARSA) Program under NIAID/NIH contract no. HHSN272200700055C. The authors thank Dr Harriet Haysom for helpful discussion of the manuscript.

## Conflicts of Interests

The authors declare no competing interests.

## Data Availability Statement

The mass spectrometry proteomics data will be deposited to the ProteomeXchange Consortium via the PRIDE partner repository upon publication.

## Notes

### Competing Interest Statement

The authors have declared no competing interest.

