## Supplementary Material for "Chemical proteomics identifies signal peptidase IB (SpsB) as a target of the SOS response inhibitor OXF-077 and a regulator of quinolone resistance emergence in *Staphylococcus aureus*"

### Supplementary Scheme

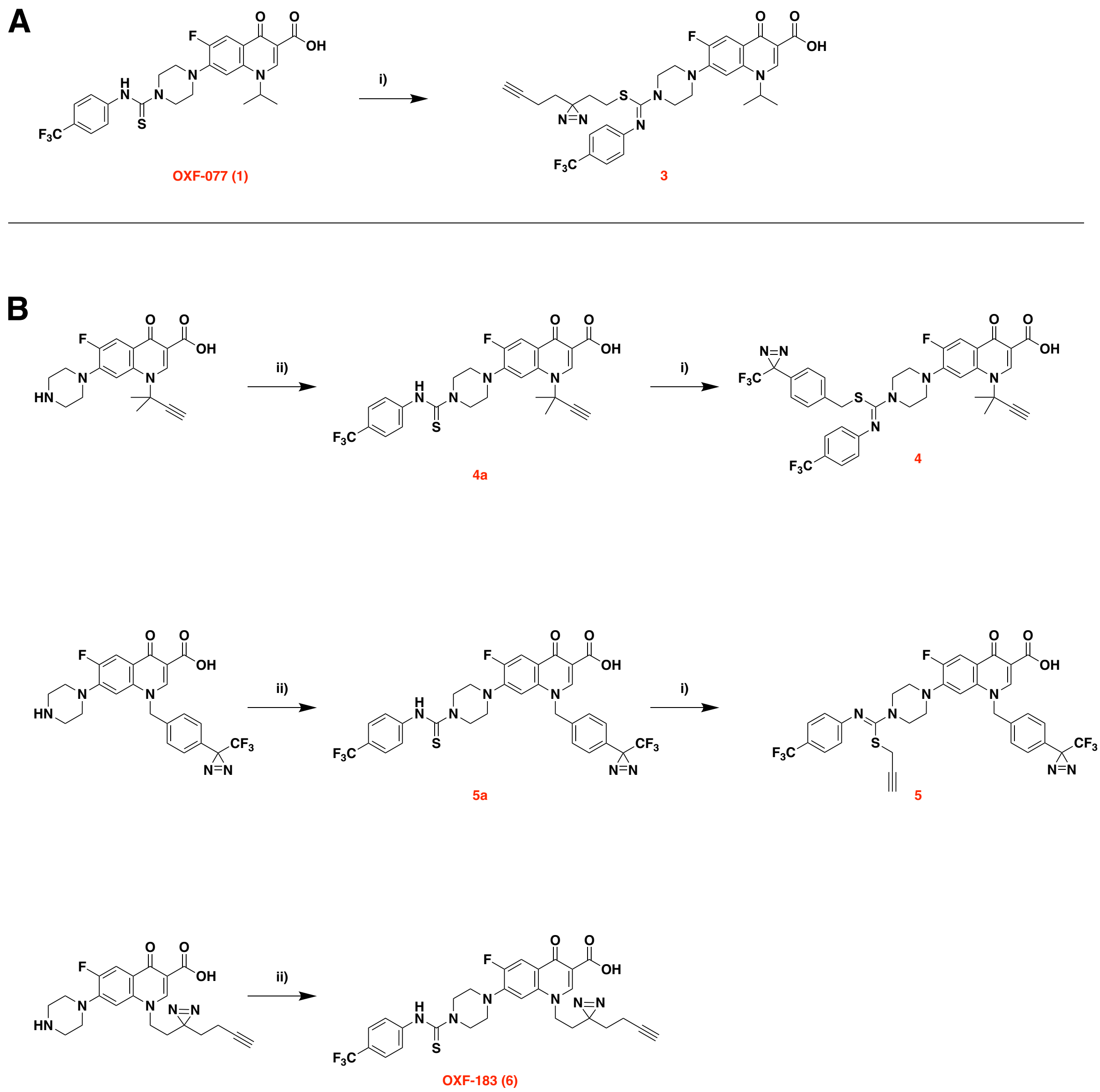

**Scheme S1 – Formation of OXF-077 Probes. (A)** Compound **3**. **(B)** Compounds **4** – **6.** General reagents and conditions: **i)** R-Br, $K_2CO_3$, MeCN, RT, 18 h. **ii)** 4-(trifluoromethyl)phenyl isothiocyanate, MeCN, 50 °C, 18 h.

### Supplementary Note

**Supplementary Note 1** – In the UniProt protein database, SpsB in JE2 *S. aureus* is annotated as being shorter than in other strains of *S. aureus* (17.6 kDa), missing the 36 amino acids N-terminal transmembrane alpha-helix. However, the UniProt sequence for SpsB in *S. aureus* COL has a ‘sequence caution’ notice that identifies that the ENA listing is incorrect due to “erroneous initiation”. Upon inspection, this DNA sequence translates identically to the JE2 peptide sequence listed on UniProt, and therefore it was assumed that the correct molecular weight of JE2 SpsB was 21.6 kDa.

### Supplementary Table

**Table S1** – Full list of all identified proteins (separate Excel file). Additionally, proteins are organised by significant enrichment (*p*-value <0.05, $log_2$(fold change) >1.5) with **OXF-183** (2 µM), significant competition (*p*-value <0.05, $log_2$(fold change) <-1) with **OXF-077** (10 and 100 µM) and both statistically significant enrichment with **OXF-183** and competition with **OXF-077** (100 µM). The mass spectrometry proteomics data will be deposited to the ProteomeXchange Consortium via the PRIDE partner repository upon publication.

### Supplementary Figures

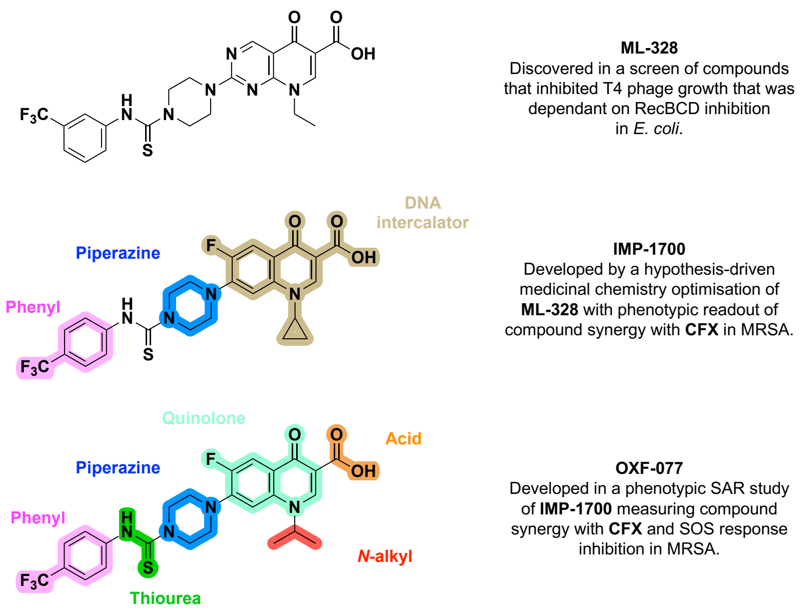

**Figure S1 – Development of ML-328, IMP-1700 and OXF-077.** Overview of prior structure–activity relationship (SAR) investigation with points of variation of **IMP-1700** and **OXF-077** colour coded as phenyl (pink), thiourea (green), piperazine (blue), quinolone (turquoise), DNA intercollator (brown), carboxylic acid (orange), and *N*-alkyl (red).

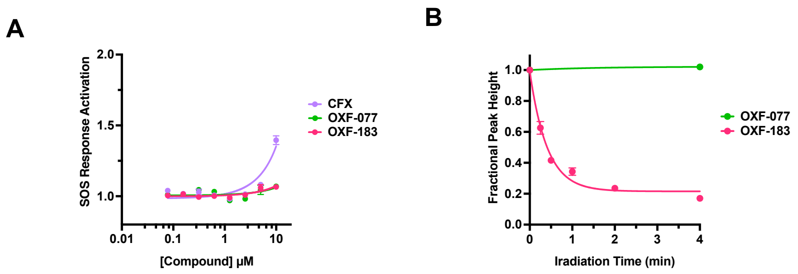

**Figure S2 – Further validation of OXF-183, a photochemical probe of OXF-077. (A)** SOS response activation (fold-change) of **OXF-183**, **OXF-077** and **CFX**. **(B)** Photoactivation of **OXF-183** determined by HPLC. Data represent mean ± SEM, n=3 independent replicates.

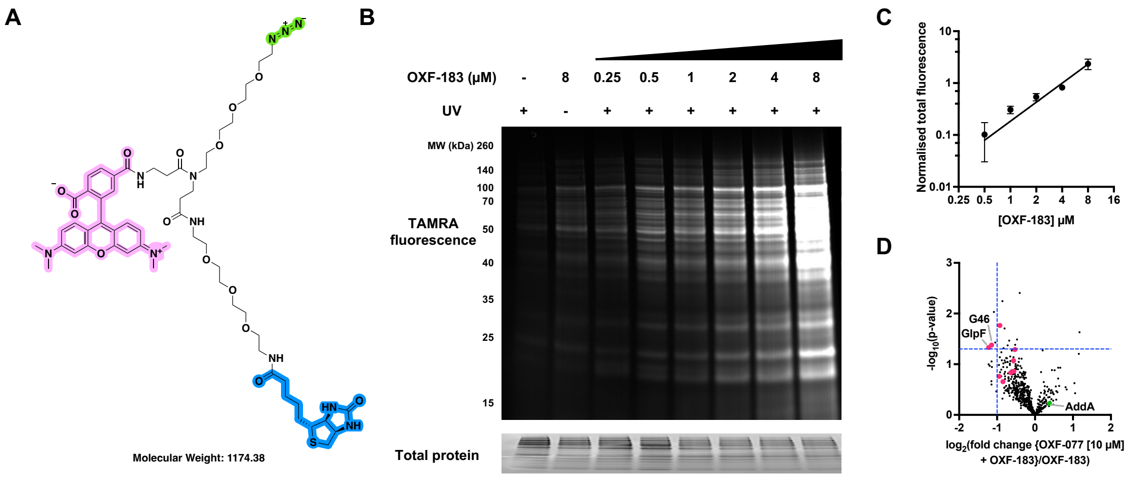

**Figure S3 – Additional affinity-based protein profiling (A*f*BPP) with OXF-077 photocrosslinking probe OXF-183.** **(A)** Structure of azide-TAMRA-biotin (AzTB), containing azide (green) for copper(I)-catalysed azide-alkyne cycloaddition (CuAAC); TAMRA (pink) for in-gel fluorescence visualisation; and biotin (blue) for enrichment of labelled proteins. **(B)** In-gel fluorescence of protein labelling in live MRSA using **OXF-0183** with or without UV and at increasing concentrations. **(C)** Quantification of the log-log relationship between fluorescence intensity and [**OXF-183**]. **(D)** Proteomic analysis of proteins enriched following **OXF-183** (2 µM) treatment in the presence or absence of **OXF-077** (10 µM) competition. Data represent mean ± standard error of the mean (SEM), n=3 independent biological replicates.

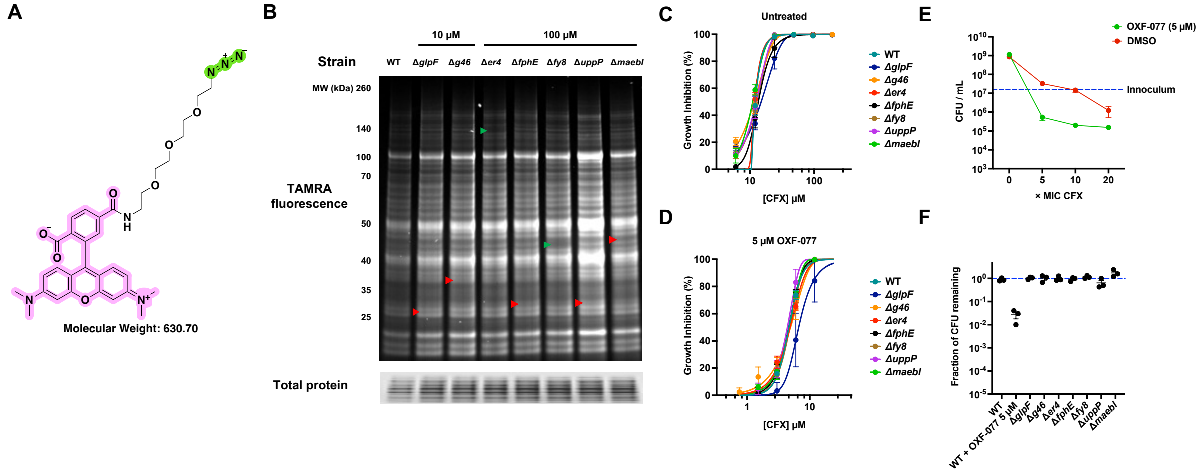

**Figure S4 – Investigation of proteins in a JE2 background.** **(A)** Structure of azide-TAMRA (AzT) containing azide (green) for copper(I)-catalysed azide-alkyne cycloaddition (CuAAC) and TAMRA (pink) for in-gel fluorescence visualisation. **(B)** In-gel A*f*BPP with **OXF-183** (2 µM) in wild-type (WT) and corresponding knockout strains of proteins identified in chemoproteomic experiments displaying competition with 10 µM and 100 µM of **OXF-077.** Bands that do (green) and do not (red) drop out are highlighted at their approximate molecular weight. **(C)** **CFX** growth inhibition (%) in untreated wild-type (WT) and JE2 mutants. **(D)** **CFX** growth inhibition (%) in **OXF-077** (5 µM) treated WT and JE2 mutants. **(E)** CFU/mL of *S. aureus* after treatment with varying concentrations of **CFX** (MIC **CFX** = 8 µg $mL^{-1}$, 24 µM) with and without **OXF-077** (5 µM). **(F)** Fraction of CFU remaining at 10 × MIC **CFX** (240 µM) in WT and JE2 mutants. Data represent mean ± standard error of the mean (SEM), n=3 independent biological replicates except in **B** where only WT and data that showed a difference to the WT (*er4* and fy8) are n=3.

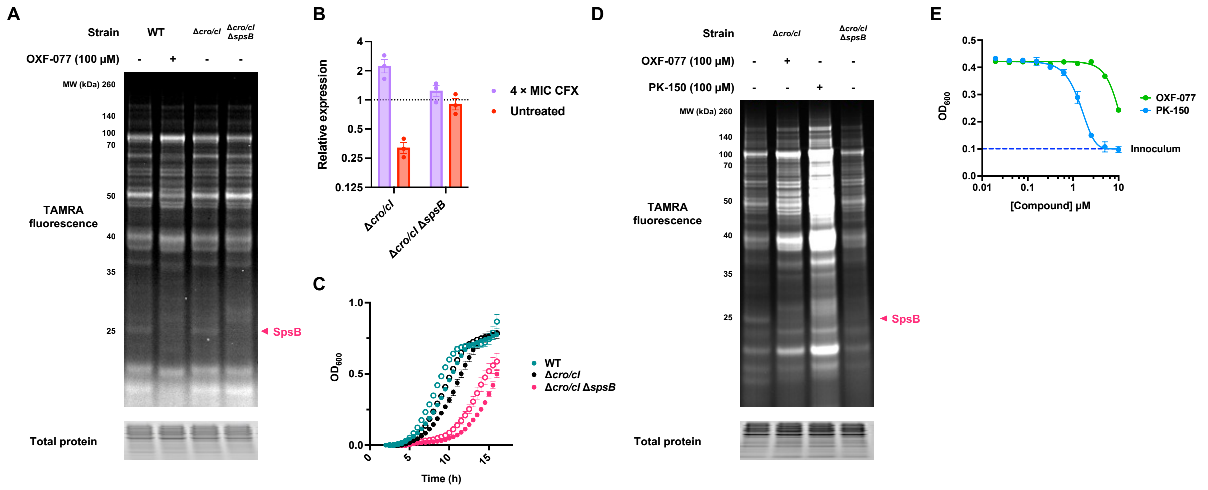

**Figure S5 – Study of SpsB activity in the RN1 strain. (A)** Uncropped in-gel *Af*BPP **OXF-183** (2 μM) in wild-type (WT) with and without competition of **OXF-077** compared with Δ*cro/cI* and Δ*cro/cI* Δ*spsB*, Fig. 3A. **(B)** Relative expression of *recA* and *rho* in Δ*cro/cI* and Δ*cro/cI* Δ*spsB* with and without treatment with 4 × MIC **CFX** (0.75 μM). **(C)** Growth time course of RN1 *S. aureus* strains in the presence (●) or absence (○) of 0.25 × MIC **CFX** (0.047 μM). **(D)** Uncropped in-gel *Af*BPP with **OXF-183** (2 μM) in Δ*cro/cI* strain with and without competition of **OXF-077** and **PK-150** compared with Δ*cro/cI* Δ*spsB*, Fig. 3F. The increased lane fluorescence of cells treated with **PK-150** possibly represents an increase in cell permeation of **OXF-183** under these treatment conditions. **(E)** Growth of *precA-gfp* JE2 *S. aureus* treated with varying concentrations of **OXF-077** or **PK-150**. Data represent mean ± standard error of the mean (SEM), n=3 independent biological replicates.

### Biological Methods

#### Bacterial strains, culture conditions and compound treatment

Bacterial strains outlined in Table S2 were revived from a frozen stock as an overnight culture grown on non-selective Muller Hinton Agar (MHA, Sigma-Aldrich, UK) supplemented with associated antibiotics at 37 °C for 17 h. Overnight cultures were grown in Muller Hinton Broth (MHB, Sigma-Aldrich, UK) supplemented with associated antibiotics to late stationary phase at 37 °C and 180 RPM in a shaking incubator (SI-200, Cole Parmer, UK). Plates were incubated at 37 °C and 180 RPM in a shaking incubator (SI-200, Cole Parmer, UK) with a BreatheEasy seal (Diversified Biotech, USA), or at 37 °C and 500 RPM a CLARIOstar Plus microplate reader (BMG, UK) without a seal. Compounds were prepared as DMSO stocks for biological experiments, except **CFX** which was dissolved in an equivalent amount of HCl. All compounds were stored at −20 °C and thawed on the day of use.

**Table S2 - Bacterial strains used in this work**

| S. aureus strain | Description | Resistance markers (Concentration) | Source |
|---|---|---|---|
| USA300 JE2 | A derivative of CA-MRSA USA300 LAC, cured of plasmids | - | Fey *et al.* 2013[1] |
| USA300 JE2 pCN34 p*recA-gfp* | JE2 containing pCN34 with GFP under the *recA* promoter | Kanamycin (50 µg/mL) | Clarke *et al.* 2019[2] |
| USA300 JE2 *glpF*::Tn*Ery* | JE2 with a transposon insertion in *glpF* | Erythromycin (10 µg/mL) | Fey *et al.* 2013[1] |
| USA300 JE2 *SAUSA300_0329*::Tn*Ery* | JE2 with a transposon insertion in *SAUSA300_1519* (*g46*) | Erythromycin (10 µg/mL) | Fey *et al.* 2013[1] |
| USA300 JE2 *SAUSA300_2213*::Tn*Ery* | JE2 with a transposon insertion in *SAUSA300_2213* (*er4*) | Erythromycin (10 µg/mL) | Fey *et al.* 2013[1] |
| USA300 JE2 *SAUSA300_2518*::Tn*Ery* | JE2 with a transposon insertion in *SAUSA300_2518* (*fphE*) | Erythromycin (10 µg/mL) | Fey *et al.* 2013[1] |
| USA300 JE2 *SAUSA300_1519*::Tn*Ery* | JE2 with a transposon insertion in *SAUSA300_0329* (*fy8*) | Erythromycin (10 µg/mL) | Fey *et al.* 2013[1] |
| USA300 JE2 *uppP*::Tn*Ery* | JE2 with a transposon insertion in *uppP* | Erythromycin (10 µg/mL) | Fey *et al.* 2013[1] |
| USA300 JE2 SAUSA300_1684::TnEry | JE2 with a transposon insertion in *SAUSA300_1684* (*maebl*) | Erythromycin (10 µg/mL) | Fey *et al.* 2013[1] |
| RN1 | NTCC 8325 RN1 *S. aureus* | - | Zhao and Bodine *et al.* 2022[3] |
| RN1 Δ*cro/cI* | RN1 with a *cro/cI* knockout | - | Zhao and Bodine *et al.* 2022[3] |
| RN1Δ*cro/cI* Δ*spsB* | RN1 with a *cro/cI* and *spsB* double knockout | - | Zhao and Bodine *et al.* 2022[3] |

#### Software analysis and plate reading

$OD_{600}$ and fluorescence intensity readings were recorded in a CLARIOstar Plus microplate reader (BMG, UK). Measurements were background corrected against a non-inoculum control of MHB and normalised to DMSO control. GraphPad Prism 10.4.2 (534) (Dotmatics, USA) was used to generate log dose-response curves for the data and calculate mean $IC_{50}$ value and SEM.

##### SOS response activation and inhibition

SOS response inhibition and activation was determined in 384-well plates (Greiner Bio-One, UK) using an SOS reporter assay. Two-fold serial dilutions of the compounds in triplicate were performed in MHB with a final volume of 25 µL and a no compound control. Overnight cultures of pCN34 *pRecA-GFP* USA300 JE2 were diluted 8-fold and supplemented with **CFX** (where stated), and 25 µL added to each well (excluding no-inoculum control) to achieve $4 \times 10^7$ CFU (final volume 50 µL, 1% (*v/v*) DMSO, and 100 µM **CFX**, where stated). Plates were incubated for 6 h after which GFP fluorescence (Ex 375, Em 425) and $OD_{600}$ were measured, and background corrected GFP/$OD_{600}$ reported.

##### Minimum inhibitory concentration (MIC)

Antibiotic susceptibility testing was determined in 384-well plates (Greiner Bio-One, UK) by MIC broth microdilution according to CLSI methods M07-A11.[4] Two-fold serial dilutions of the compounds in triplicate were performed in MHB with a final volume of 25 µL and a no compound growth control. A direct colony suspension was made by dispersing singular, well isolated bacterial colonies from the overnight revive plates in 3 mL sterile Phosphate-Buffered Saline (PBS) to achieve a turbidity of 0.5 McFarland standard (Oxoid, UK), approximately $10^8$ CFU/ml. The inoculum was vortexed and then further diluted 1:100 in MHB to achieve a final inoculum of $10^6$ CFU/mL. Inoculum (25 µL) containing a fixed concentration of a second compound (or DMSO) was added to each well to achieve a final CFU/mL of $5 \times 10^5$, excluding no-inoculum sterility control which had only MHB added (final volume 50 µL, 1% (*v/v*) DMSO). Plates were incubated overnight for 16 - 18 h after which the $OD_{600}$ was recorded, or read kinetically overnight with $OD_{600}$ recordings every 30 min.

##### Photoactivated degradation

Compounds were made up to 50 µM in 250 µl PBS in Eppendorf tubes. Samples were irradiated on ice for the designated time in a UV Irradiator (Wavey Technologies, UK). Samples were kept on ice and in the dark when not irradiated. Samples were then analysed by High Performance Liquid Chromatography (HPLC) analysis using an SPD-20A UV detector (Shimadzu, Japan) set to 280 nm and an ACE Equivalence 3, C18, 150 × 4.6 mm column (Avantor, USA), with a sample injection volume of 25 µL.

##### Affinity based protein profiling (A*f*BPP)

Overnight cultures were pelleted (35000 × g, 10 min, 4 °C), washed twice in PBS, and resuspended in an equivalent volume of PBS. 1 mL aliquots of resuspension were incubated (500 rpm, 1 h, 37 °C) with required compound(s) or DMSO. Samples were UV-irradiated on ice for 1 min, where stated. Samples were centrifuged (17000 × g, 5 min, 4 °C), the supernatant discarded, and pellets were resuspended in 300 µL lysis buffer (1% (v/v) Triton X-100, 1% (*w/v*) sodium dodecyl sulfate (SDS), EDTA-free complete protease inhibitor cocktail (1 ×, Roche, Switzerland) in PBS). Samples were added to Pathogen Lysis tubes (S, Quiagen, Germany) and vortexed for 15 min. Samples were then centrifuged (17000 × g, 5 mins, 4 °C) and the supernatant placed into 2 mL Eppendorf tubes and boiled at 95 °C for 10 minutes to attenuate. Supernatant protein concentration was determined using the DC Protein Assay (BioRad, USA) as per manufacturer's instructions. Samples were adjusted to 1 mg/mL in PBS to 100 µL.

##### CuACC ligation

AzTB or AzT (0.1 mM), $CuSO_4$ (1 mM), TBTA (0.1 mM), TCEP (1 mM) were premixed, and added to 100 µL lysate. Reaction mixtures were incubated (500 rpm, 1 hr, RT), then quenched (5 mM EDTA) on ice. Protein was precipitated by sequential addition of MeOH (200 µL), $CHCl_3$ (50 µL) and $H_2O$ (100 µL) to each sample at on ice. Samples were vortexed and pelleted (17000 × g, 5 min, 4 °C). The top layer (MeOH/$H_2O$) was removed and 300 µL MeOH added, the samples were sonicated (5 mins), pelleted (17000 × g, 5 min, 4 °C) and the resulting pellet washed with MeOH (300 µL × 3). Pellets were air dried (5 min), resuspended in 20 µL of 1% SDS in PBS and probe sonicated (10% amplitude, 10 s) on ice, then made up to 1 mg/mL by adding 80 µL of PBS.

##### In-gel fluorescence

Gels were electrophoresed using an Invitrogen XCell SureLock Electrophoresis Mini-Cell Tank with an Invitrogen PowerEase Touch 350 W Power Supply (Thermo Fisher Scientific, USA). SDS-PAGE samples were prepared by sequential addition of 5 µL 4 × NuPAGE LDS Sample Buffer (Thermo Fisher Scientific, USA), 0.5 M DTT (2 µL) and 13 µL of protein sample after CuACC ligation. Samples were boiled at 5 min at 95 °C, loaded onto 12% Bis-Tris gels and separated by SDS-PAGE. TAMRA fluorescence was imaged at 520 nm excitation on an Odyssey M (LI-COR, USA). Total protein was imaged after staining with QuickBlue Protein Stain (Stratech Scientific Ltd, UK) at 700 nm excitation on an Odyssey M.

##### Pull-down, on bead peptide digest, elution and tandem LC-MS/MS

CuAAC ligation and protein precipitation was performed as described previously with the reaction scaled up to 1 mg of each condition, protein concentrations adjusted to 2.5 mg/mL and cells lysed in a mechanical bead beater for 8 min at 8.5 M/s (MP Biomedicals, USA). 100 µL of 50% slurry NeutrAvidin Agarose Resin (Thermo Fisher Scientific, USA) per 1 mg of protein was prewashed (3000 × g, 2 mins, 4 °C) three times with excess 0.2% SDS in PBS. Beads were aliquoted and incubated with sample (1200 rpm, 2 h, RT). Samples were centrifuged, supernatant was removed, and beads washed three times as before. Beads were washed in Urea AmBic buffer (8 M Urea in 100 mM AmBic buffer, pH 7.8, 2 × 100 µL), resuspended in 1.5 × volume Urea AmBic buffer and incubated (700 rpm, 10 min, RT). TCEP was added (10 mM) and incubated (30 min, RT). 2-chloroacetamide was added (50 mM) and incubated (30 min, RT). Samples were pre-digested using 1 µg LysC per 100 µg protein (700 rpm, 2 h, 37 °C). Urea was diluted to 2 M in 100 mM AmBic, $CaCl_2$ was added (2 mM), and 1 µg trypsin per 40 µg protein was added before incubation (700 rpm, 16 h, 37 °C). 5% formic acid was used to quench the reaction. Samples were centrifuged (17000 × g, 30 min, 4 °C), supernatant desalted on C18 columns, eluted with 50% acetonitrile containing 0.1% TFA and dried.

Peptides were separated by nano liquid chromatography (Thermo Scientific Ultimate 3000 RSLC) coupled to a Q Exactive mass spectrometer equipped with an Easy-Spray source (Thermo Fisher Scientific, USA). Peptides were trapped onto a C18 PepMap100 precolumn (5 µm, 100 Å, 300 µm × 5 mm, Thermo Fisher Scientific, USA) using Solvent A (0.1% formic acid, HPLC grade water). The peptides were further separated onto an Easy-Spray C18 column (2 µm, 100 Å, 75 µm × 50 cm, Thermo Fisher Scientific, USA) using a 60 min linear gradient (15% to 35% solvent B (0.1% formic acid in acetonitrile)) at a flow rate 200 nL $min^{-1}$. The raw data were acquired on the mass spectrometer in a data-dependent acquisition mode (DDA). Full-scan MS spectra were acquired in the Orbitrap (Scan range 350-1500 m/z, resolution 70,000; AGC target, $3 \times e^6$, maximum injection time, 50 ms). The 10 most intense peaks were selected for higher-energy collision dissociation (HCD) fragmentation at 30% of normalized collision energy. HCD spectra were acquired in the Orbitrap at resolution 17,500, AGC target $5 \times e^4$, maximum injection time 120 ms with fixed mass at 180 m/z. Charge exclusion was selected for unassigned and 1+ ions. The dynamic exclusion was set to 20 s.

**Chemoproteomics data analysis**

Tandem mass spectra were searched using Sequest HT in Proteome Discoverer version 1.4 against a *Staphylococcus aureus* (strain USA300) protein sequence database containing 2607 protein entries (Proteome ID: UP000001939) and 292 common laboratory contaminants. Carbamidomethylation (C) was selected as a fixed modification. Oxidation (M), N-terminal acetylation, Carbamylation (K) and Deamidation (N, Q) were selected as variable modifications. Trypsin was chosen as the endoprotease. Two missed cleavages were permitted. Peptide mass tolerance was set at 20 ppm on the precursor and 0.6 Da on the fragment ions. Data was filtered at FDR below 1% at the peptide level. Downstream bioinformatic analysis was performed using Perseus 2.0.11.0.[5] Median normalization was performed on the identified hits followed by filtering data that do not have 67% valid values amongst the 3 replicates. Imputation was carried out by replacing the missing values from the normal distribution. An unpaired Student's T-test was performed between the tested conditions at $S_0 = 2$ and $p$-value of ≤0.05.

**Persister cell formation**

Adapted from Johnson *et al.*, 2013.[6] Overnight cultures were diluted 100-fold in MHB and CFU/mL assessed. Cultures split into 1 mL in Eppendorf's and challenged with increasing concentrations of **CFX** (5, 10, 20 × MIC), with or without **OXF-077** (5 µM). To enumerate survivors, 10 µL culture was taken, serially diluted, plated on MH agar plates and colonies were counted after 24 h of growth.

**Frequency of resistance (FoR) to CFX**

Overnight cultures of RN1 *S. aureus* were diluted 5-fold in MHB and CFU/mL assessed. 250 µL deposited MH agar plates containing concentrations at 4 × MIC **CFX** (0.75 µM). After 24, 48 and 72 h, the number of colonies were counted for each condition and compared to the initial CFU/mL to assess FoR.

##### *recA* expression upon CFX treatment

Adapted from Cirz *et al.* 2006.[7] 10 µL of overnight cultures of RN1 *S. aureus* were added to 1 mL of MHB and grown for 3 h. Cultures were then split into 0.5 mL aliquots, of which one was treated with 4 × MIC of **CFX** (0.75 µM) for 2 h, after which cultures were added to 1 mL of RNAprotect (Quiagen, Germany), vortexed, incubated at RT (5 min) and pelleted (5,000 × g, 10 min, 4 °C), supernatant removed and pellets stored at −80 °C.

RNA was extracted with a RNeasy Mini Kit (Quiagen, Germany) and performed by manufacturer's instructions during which lysis was achieved by mechanical lysis in pathogen lysis tubes (Quiagen, Germany). RNA extraction included on collumn DNA digestion using a RNase-Free DNAse Set (Quiagen, Germany). cDNA was synthesised using a QuantiTect Reverse Transcription Kit (Quiagen, Germany) and performed by manufacturer's instructions.

qPCR was performed as described by Sihto *et al.* 2014,[8] using primers (250 nM) outlined in Table S3 monitored using the SYBR Green fluorogenic stain (BioRad, USA). Reactions were cycled in CFX Opus 96 Real-Time PCR system (BioRad, USA) with the following cycling conditions: hot start (8 min, 95 °C), 45 amplification cycles (95 °C for 10 s, 57 °C for 15 s, 72 °C for 20 s, 78 °C for 1 s with a single fluorescence measurement), a melting curve (60–95 °C at 0.5 °C per cycle with 5 s dwell, continuous fluorescence measurement), and a final cooling step.

qPCR data was extracted with CFX Maestro (BioRad, USA) and *recA* Cq values normalised to *rho* Cq values (ΔCq). Resulting ΔCq values of treated and untreated cells were compared (ΔΔCq). Data was represented as relative expression (= $2^{-\Delta Cq}$) and fold-change in expression (= $2^{-\Delta\Delta Cq}$). Results are representative of 3 independent biological replicates.

**Table S3 - Primers used in this work**

| Gene | Forward Primer | Reverse Primer | Source |
|---|---|---|---|
| *recA* | 5'-AAGTACGTCGTGCAGA-3' | 5'-TGACCCATTCGTTCGC-3' | Cirz *et al.* 2006.[7] |
| *rho* | 5'-GAAGCTGCTGAAGTCG-3' | 5'-CGTCCATACGTGAACCC-3' | Cirz *et al.* 2006.[7] |

### Chemical Methods

#### General information

Materials were purchased from commercial suppliers and used as received. Analytical thin-layer chromatography (TLC) was performed on 0.25 mm silica gel 60 F254 pre-coated plates 0.25 mm (Merck, UK) and visualized under ultraviolet light (254 and 365 nm). Purification by column chromatography was carried out using a CombiFlash $R_f$ automated column system with RediSep silver disposable flash columns (Teledyne, USA).

$^{1}$H nuclear magnetic resonance (NMR) and $^{13}$C NMR spectra were recorded at room temperature at 400 MHz and 101 MHz respectively (Bruker, USA). Chemical shifts are reported as parts per million (δ). The spectra are calibrated using the solvent peak with the data provided by Fulmer *et al.*[9] The multiplicity of each signal is indicated by; s (singlet); br s (broad singlet); d (doublet); t (triplet); q (quartet); p (pentet); hept (heptet); m (multiplet) or combinations thereof. Coupling constants (*J*) are reported in hertz (Hz) and averaged for interacting protons. Low-resolution mass spectroscopy (LRMS) and high-resolution mass spectroscopy (HRMS) was collected on a BioAccord (Waters, USA). HPLC analysis was conducted using an SPD-20A UV detector (Shimadzu, Japan) with 254 nm and 280 nm detection, and an ACE Equivalence 3, C18, 150 × 4.6 mm column (Avantor, USA).

#### Synthetic procedures

**7-(4-(((2-(3-(but-3-yn-1-yl)-3H-diazirin-3-yl)ethyl)thio)((4-(trifluoromethyl)phenyl)imino)methyl)piperazin-1-yl)-6-fluoro-1-isopropyl-4-oxo-1,4-dihydroquinoline-3-carboxylic acid** (**3**)

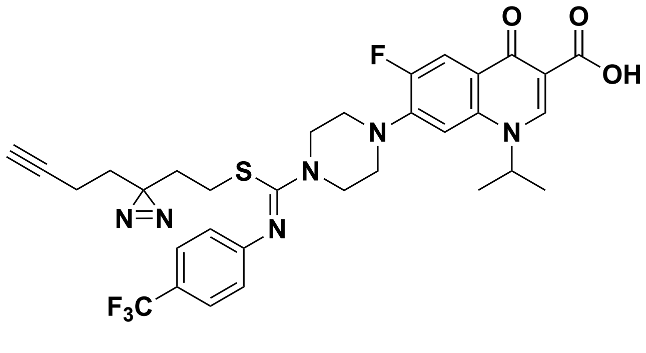

6-Fluoro-1-isopropyl-4-oxo-7-(4-((4-(trifluoromethyl)phenyl)carbamothioyl)piperazin-1-yl)-1,4-dihydroquinoline-3-carboxylic acid (**2**, **OXF-077**[10]) (25 mg, 0.046 mmol) and $K_2CO_3$ (13 mg, 0.093 mmol) were suspended in anhydrous MeCN (1 mL) and 3-(but-3-yn-1-yl)-3-(2-iodoethyl)-3H-diazirine (23 mg, 0.093 mmol) was added. The reaction was stirred at RT for 36 h after which sat. aq. $NH_4Cl$ (10 mL) was added, and the suspension was stirred for 10 min. The product was extracted with DCM (3 × 5 mL), dried ($MgSO_4$), and purified by flash column chromatography, 0–5% MeOH in DCM to afford a colourless solid (20 mg, 66%). $R_f$ = 0.77 ($SiO_2$; DCM:MeOH, 9:1); **$^1$H NMR** (400 MHz, $CDCl_3$) δ 15.08 (s, 1H), 8.80 (s, 1H), 8.10 (d, *J* = 12.9 Hz, 1H), 7.54 (d, *J* = 8.3 Hz, 2H), 7.02 (d, *J* = 6.8 Hz, 1H), 6.97 (d, *J* = 8.3 Hz, 2H), 4.93 (p, *J* = 6.7 Hz, 1H), 3.92 (t, *J* = 5.0 Hz, 4H), 3.39 (t, *J* = 5.0 Hz, 4H), 2.26 (t, *J* = 7.5 Hz, 2H), 1.99–1.92 (m, 3H), 1.68 (d, *J* = 6.7 Hz, 6H), 1.56 (dt, *J* = 24.0, 7.5 Hz, 4H); **$^{13}$C NMR** (101 MHz, $CDCl_3$) δ 176.62 , 167.22, 154.60 (d, *J* = 17.0 Hz), 152.18, 145.62 (d, *J* = 10.6 Hz), 142.97, 137.57, 126.17 (q, *J* = 4.0 Hz, 2C), 125.91, 124.47 (q, *J* = 32.4 Hz), 121.39 (2C), 113.03 (d, *J* = 23.0 Hz), 108.50, 103.89, 82.36, 77.24, 69.46, 52.64, 49.79 (d, *J* = 4.6 Hz, 2C), 47.80 (2C), 33.29, 31.90, 27.31, 26.68, 22.23, 13.20; **$^{19}$F NMR** (376 MHz, $CDCl_3$) δ: -61.61, -120.20; **LRMS** m/z (ESI$^+$) 657 ([M+H]$^+$); **HRMS** m/z (ESI$^+$) found 657.2278, $C_{32}H_{33}F_4N_6O_3S$ ([M+H]$^+$) requires 657.2266; **HPLC** Retention time 13.8 min, 98% (280 nm).

**6-fluoro-1-(2-methylbut-3-yn-2-yl)-4-oxo-7-(4-((4-(trifluoromethyl)phenyl)carbamothioyl)piperazin-1-yl)-1,4-dihydroquinoline-3-carboxylic acid** (**4a**)

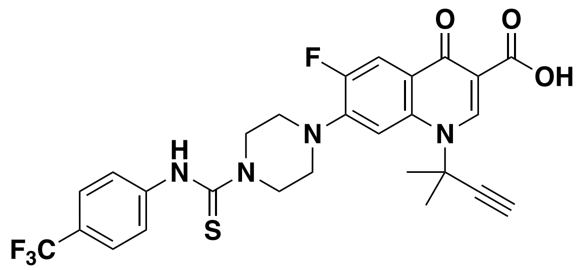

6-Fluoro-1-(2-methylbut-3-yn-2-yl)-4-oxo-7-(piperazin-1-yl)-1,4-dihydroquinoline-3-carboxylic acid[11] (50 mg, 0.14 mmol) and 4-(trifluoromethyl)phenyl isothiocyanate (56 mg, 0.28 mmol) were suspended in 2 mL of anhydrous MeCN. The reaction mixture was stirred for 4 h at RT after which the solvent was removed *in vacuo*. The solid was purified by flash column chromatography, 0–5% MeOH in DCM to give a colourless solid (79 mg, 84%). $R_f$ = 0.70 ($SiO_2$; DCM:MeOH, 90:10); **$^1$H NMR** (400 MHz, $CDCl_3$) δ 8.87 (s, 1H), 8.23 (s, 1H), 8.02–7.94 (m, 2H), 7.56 (d, *J* = 8.4 Hz, 2H), 7.44 (d, *J* = 8.4 Hz, 2H), 5.30 (s, 2H), 4.21 (t, *J* = 5.3 Hz, 4H), 3.48 (t, *J* = 5.1 Hz, 4H), 2.98 (s, 1H), 2.10 (s, 6H); **$^{13}$C NMR** (101 MHz, $CDCl_3$) δ 182.80, 176.72, 167.50, 152.90 (d, *J* = 251.8 Hz), 143.60 (d, *J* = 10.2 Hz), 143.38, 143.08, 136.79, 126.93, 126.61, 126.15 (q, *J* = 5.7 Hz), 123.08, 121.85, 112.76 (d, *J* = 23.3 Hz), 108.91, 107.67, 83.51, 77.27, 58.80, 53.47, 49.12, 49.07, 48.67, 30.78; **$^{19}$F NMR** (376 MHz, DMSO-$d_6$) δ: -61.69, -121.43; **LRMS** m/z (ESI$^+$) 561 ([M+H]$^+$); **HRMS** m/z (ESI$^+$) found 561.1579, $C_{27}H_{25}F_4N_4O_3S$ ([M+H]$^+$) requires 561.1578.

**6-fluoro-1-(2-methylbut-3-yn-2-yl)-4-oxo-7-(4-((4-(3-(trifluoromethyl)-3*H*-diazirin-3-yl)benzyl)(4-(trifluoromethyl)phenyl)carbamothioyl)piperazin-1-yl)-1,4-dihydroquinoline-3-carboxylic acid** (**4**)

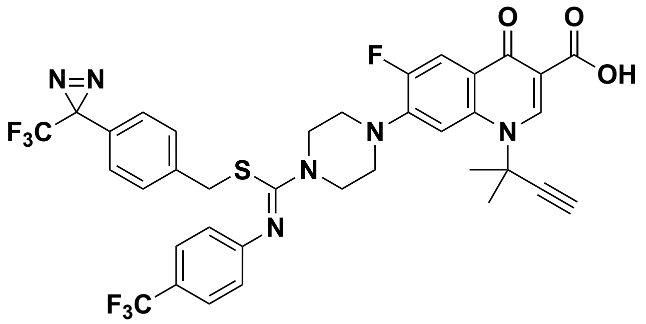

6-Fluoro-1-(2-methylbut-3-yn-2-yl)-4-oxo-7-(4-((4-(trifluoromethyl)phenyl)carbamothioyl)piperazin-1-yl)-1,4-dihydroquinoline-3-carboxylic acid (**4a**) (30 mg, 0.054 mmol) and $K_2CO_3$ (7.4 mg, 0.054 mmol) were suspended in anhydrous MeCN (1 mL) and 4-[3-(trifluoromethyl)-3H-diazirin-3-yl]benzyl bromide (15 mg, 0.054 mmol) was added. The reaction was stirred at RT for 24 h after which sat. aq. $NH_4Cl$ (5 mL) and $H_2O$ (5 mL) were added, and the suspension was stirred for 10 min. The product was extracted with DCM (3 × 5 mL), dried ($MgSO_4$), and purified by flash column chromatography, 0–5% MeOH in DCM to afford a colourless crystalline solid (23 mg, 55%). $R_f$ = 0.51 ($SiO_2$; DCM:MeOH, 9:1); **$^1$H NMR** (400 MHz, $CDCl_3$) δ 14.93 (s, 1H), 8.96 (s, 1H), 8.10 (d, *J* = 12.9 Hz, 1H), 7.96 (d, *J* = 7.0 Hz, 1H), 7.51 (d, *J* = 8.2 Hz, 2H), 7.17 (dd, *J* = 11.9, 8.5 Hz, 2H), 6.89 (d, *J* = 8.1 Hz, 1H), 3.85 (t, *J* = 5.0 Hz, 5H), 3.64 (s, 2H), 3.30 (t, *J* = 4.8 Hz, 4H), 2.88 (s, 1H), 2.15 (s, 7H); **$^{13}$C NMR** (101 MHz, $CDCl_3$) δ 176.78 (d, *J* = 1.9 Hz), 167.17, 154.39, 153.83, 152.20, 151.88, 143.92 (d, *J* = 10.0 Hz), 143.35, 139.15, 136.77, 129.12, 128.52, 126.77, 126.05 (q, *J* = 3.5 Hz), 124.69, 124.37, 122.07 (d, *J* = 7.7 Hz), 121.66, 120.64, 112.84 (d, *J* = 23.0 Hz), 109.08 (d, *J* = 2.1 Hz), 107.96, 83.74, 58.61, 49.68 (d, *J* = 4.7 Hz), 47.60, 36.20, 30.85; **$^{19}$F NMR** (376 MHz, $CDCl_3$) δ: -61.61, -65.73, -121.20; **LRMS** m/z (ESI$^+$) 759 ([M+H]$^+$); **HRMS** m/z (ESI$^+$) found 759.1985, $C_{36}H_{30}F_7N_6O_3S$ ([M+H]$^+$) requires 759.1983; **HPLC** Retention time 15.1 min, 99% (280 nm).

**6-fluoro-4-oxo-1-(4-(3-(trifluoromethyl)-3*H*-diazirin-3-yl)benzyl)-7-(4-((4-(trifluoromethyl)phenyl)carbamothioyl)piperazin-1-yl)-1,4-dihydroquinoline-3-carboxylic acid** (**5a**)

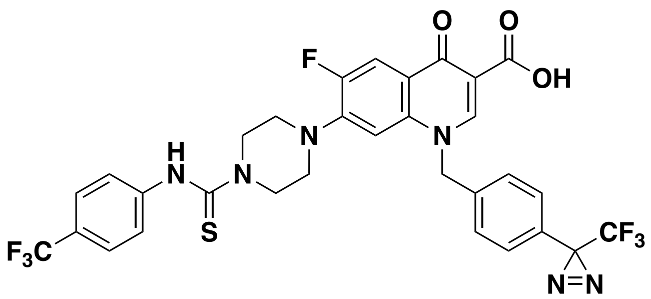

6-Fluoro-4-oxo-7-(piperazin-1-yl)-1-(4-(3-(trifluoromethyl)-3H-diazirin-3-yl)benzyl)-1,4-dihydroquinoline-3-carboxylic acid[11] (70 mg, 0.14 mmol) and 4-(trifluoromethyl)phenyl isothiocyanate (58 mg, 0.28 mmol) were suspended in 2 mL of anhydrous MeCN. The reaction mixture was stirred for 18 h after which the solvent was removed *in vacuo*. The solid was purified by flash column chromatography, 0–5% MeOH in DCM to give a colourless solid (66 mg, 66%). $R_f$ = 0.45 ($SiO_2$; DCM:MeOH, 90:10); **$^1$H NMR** (400 MHz, DMSO-*d*$_6$) δ 15.23 (s, 1H), 9.71 (s, 1H), 9.21 (s, 1H), 7.93 (d, *J* = 13.3 Hz, 1H), 7.66 (d, *J* = 8.5 Hz, 2H), 7.58 (d, *J* = 8.5 Hz, 2H), 7.57 (d, *J* = 8.1 Hz, 2H), 7.31 (d, *J* = 8.1 Hz, 2H), 7.04 (d, *J* = 7.2 Hz, 1H), 5.92 (s, 2H), 4.07 (br s, 4H), 3.33 (br s, 4H); **$^{13}$C NMR** (101 MHz, DMSO-*d*$_6$) δ 181.79, 176.89 (d, *J* = 2.7 Hz), 166.49, 152.91 (d, *J* = 249.9 Hz), 150.19, 145.24, 144.90 (d, *J* = 10.3 Hz), 138.30, 137.94, 128.40 (2C), 127.85, 127.59 (2C), 125.63 (q, *J* = 3.9 Hz), 124.69 (4C), 124.34 (q, *J* = 31.8 Hz), 123.62 (q, *J* = 7.1 Hz), 120.89, 119.71 (d, *J* = 7.4 Hz), 111.85 (d, *J* = 23.1 Hz), 107.84, 106.60, 79.63, 56.36, 49.07 (2C), 48.04 (2C), 28.43 (q, *J* = 40.1 Hz); **$^{19}$F NMR** (376 MHz, DMSO-*d*$_6$) δ: -60.37, -64.66, -121.58; **LRMS** m/z (ESI$^+$) 693 ([M+H]$^+$); **HRMS** m/z (ESI$^+$) found 693.1524, $C_{31}H_{24}F_7N_6O_3S$ ([M+H]$^+$) requires 693.1513.

**6-Fluoro-4-oxo-7-(4-((prop-2-yn-1-ylthio)((4-(trifluoromethyl)phenyl)imino)methyl)piperazin-1-yl)-1-(4-(3-(trifluoromethyl)-3*H*-diazirin-3-yl)benzyl)-1,4-dihydroquinoline-3-carboxylic acid** (**5**)

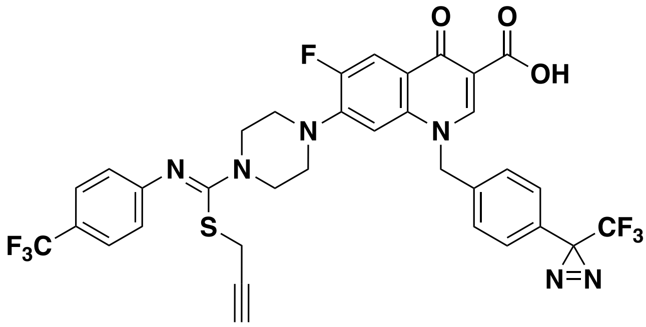

6-Fluoro-4-oxo-1-(4-(3-(trifluoromethyl)-3*H*-diazirin-3-yl)benzyl)-7-(4-((4-(trifluoromethyl)phenyl)carbamothioyl)piperazin-1-yl)-1,4-dihydroquinoline-3-carboxylic acid (**5a**) (40 mg, 0.057 mmol) and $K_2CO_3$ (12 mg, 0.87 mmol) were suspended in anhydrous MeCN (2 mL) and propargyl bromide (10 mg, 0.87 mmol) was added. The reaction was stirred at RT for 24 h after which sat. aq. $NH_4Cl$ (10 mL) was added, and the suspension was stirred for 10 min. The product was extracted with DCM (3 × 5 mL), dried ($MgSO_4$), and purified by flash column chromatography, 0–5% MeOH in DCM to afford a colourless solid (23 mg, 53%). $R_f$ = 0.63 ($SiO_2$; DCM:MeOH, 9:1); **$^1$H NMR** (400 MHz, $CDCl_3$) δ 14.90 (s, 1H), 8.78 (s, 1H), 8.05 (d, *J* = 12.9 Hz, 1H), 7.53 (d, *J* = 8.4 Hz, 2H), 7.23 (s, 4H), 7.01 (d, *J* = 8.4 Hz, 2H), 6.62 (d, *J* = 6.8 Hz, 1H), 5.48 (s, 2H), 3.85 (t, *J* = 5.0 Hz, 4H), 3.20 (t, *J* = 5.0 Hz, 4H), 3.12 (d, *J* = 2.6 Hz, 2H); **$^{13}$C NMR** (101 MHz, $CDCl_3$) δ 177.20, 166.81, 153.30 (d, *J* = 273.0 Hz) 152.59 (d, *J* = 129.6 Hz), 148.51, 145.54 (d, *J* = 10.5 Hz), 137.33, 135.27, 130.31, 127.72 (4C), 126.67 (2C), 126.12 (q, *J* = 3.7 Hz), 124.73 (d, *J* = 32.3 Hz), 123.22 (q, *J* = 3.0 Hz), 121.52 (2C), 120.78 (d *J* = 8.1 Hz), 120.50, 112.97 (d, *J* = 23.4 Hz), 108.68, 104.96, 78.96, 77.24, 72.05, 57.89, 49.41 (d, *J* = 5.0 Hz, 2C), 47.76 (2C), 20.53[1]; **$^{19}$F NMR** (376 MHz, $CDCl_3$) δ: -61.66, -65.12, -120.14; **LRMS** m/z (ESI$^+$) 731 ([M+H]$^+$); **HRMS** m/z (ESI$^+$) found 731.1678, $C_{34}H_{22}F_7N_6O_3S$ ([M+H]$^+$) requires 731.1670; **HPLC** Retention time 14.7 min, 95% (280 nm).

[1] No signal observed for C-diazirene, ~25 ppm, due to fluorine coupling.

**1-(2-(3-(but-3-yn-1-yl)-3H-diazirin-3-yl)ethyl)-6-fluoro-4-oxo-7-(4-((4-(trifluoromethyl)phenyl)carbamothioyl)piperazin-1-yl)-1,4-dihydroquinoline-3-carboxylic acid** (**6, OXF-183**)

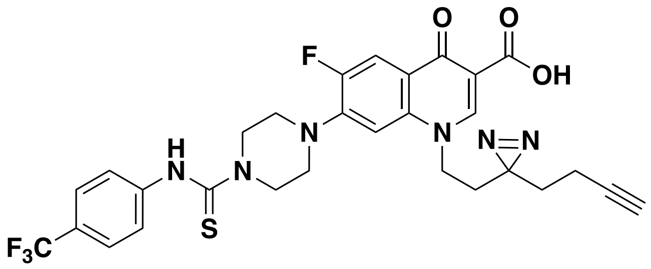

1-(2-(3-(But-3-yn-1-yl)-3H-diazirin-3-yl)ethyl)-6-fluoro-4-oxo-7-(piperazin-1-yl)-1,4-dihydroquinoline-3-carboxylic acid[11] (**6a**) (33 mg, 0.080 mmol) and 4-(trifluoromethyl)phenyl isothiocyanate (32 mg, 0.16 mmol) were suspended in anhydrous MeCN (2 mL). The reaction mixture was stirred for 4 h at RT after which the solvent was removed *in vacuo*. The solid was purified by flash column chromatography, 0–5% MeOH in DCM to give a colourless solid (39 mg, 79%). $R_f$ = 0.74 ($SiO_2$; DCM:MeOH, 90:10); **$^1$H NMR** (400 MHz, DMSO-*d$_6$*) δ 15.26 (s, 1H), 9.73 (s, 1H), 9.08 (s, 1H), 7.96 (d, *J* = 13.2 Hz, 1H), 7.67 (d, *J* = 8.6 Hz, 2H), 7.59 (d, *J* = 8.6 Hz, 2H), 7.03 (d, *J* = 7.3 Hz, 1H), 4.62 (t, *J* = 7.4 Hz, 2H), 4.16 (t, *J* = 5.1 Hz, 4H), 3.50 (t, *J* = 5.1 Hz, 4H), 2.81 (t, *J* = 2.6 Hz, 1H), 2.05–1.92 (m, 4H), 1.70 (t, *J* = 7.4 Hz, 2H); **$^{13}$C NMR** (101 MHz, DMSO-*d$_6$*) δ 181.69, 176.77, 166.47, 153.09 (d, *J* = 249.1 Hz), 149.85, 145.25, 137.49, 125.65 (q, *J* = 3.8 Hz), 124.74, 124.49, 119.70, 111.81 (d, *J* = 22.6 Hz), 107.67, 106.11, 83.55, 72.35, 49.15, 48.23, 40.47, 32.12, 31.23, 27.17, 13.04; **$^{19}$F NMR** (376 MHz, DMSO-*d$_6$*) δ: -60.33, -121.59; **LRMS** m/z (ESI$^+$) 615 ([M+H]$^+$); **HRMS** m/z (ESI$^+$) found 615.1809, $C_{29}H_{27}F_4N_6O_3S$ ([M+H]$^+$) requires 615.1796; **HPLC** Retention time 12.5 min, 97% (280 nm).

### NMR Spectra of Tested Compounds

#### 7-(4-(((2-(3-(but-3-yn-1-yl)-3H-diazirin-3-yl)ethyl)thio)((4-(trifluoromethyl)phenyl)imino)methyl)piperazin-1-yl)-6-fluoro-1-isopropyl-4-oxo-1,4-dihydroquinoline-3-carboxylic acid (3)

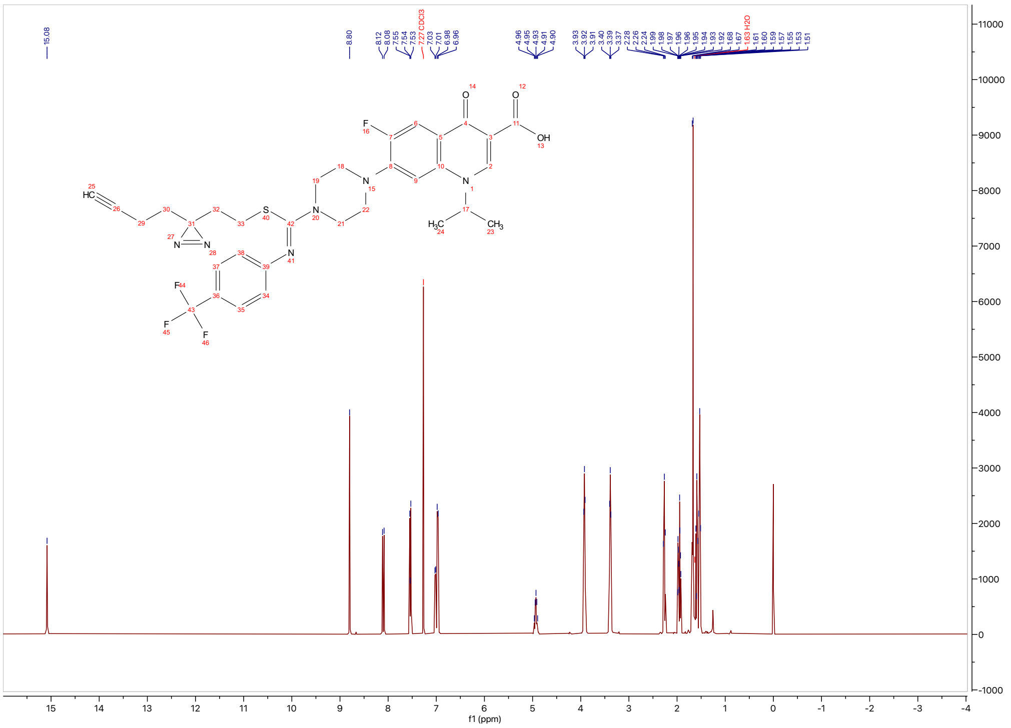

$^{1}H$ NMR (400 MHz, $CDCl_3$) at 298 K of **3**.

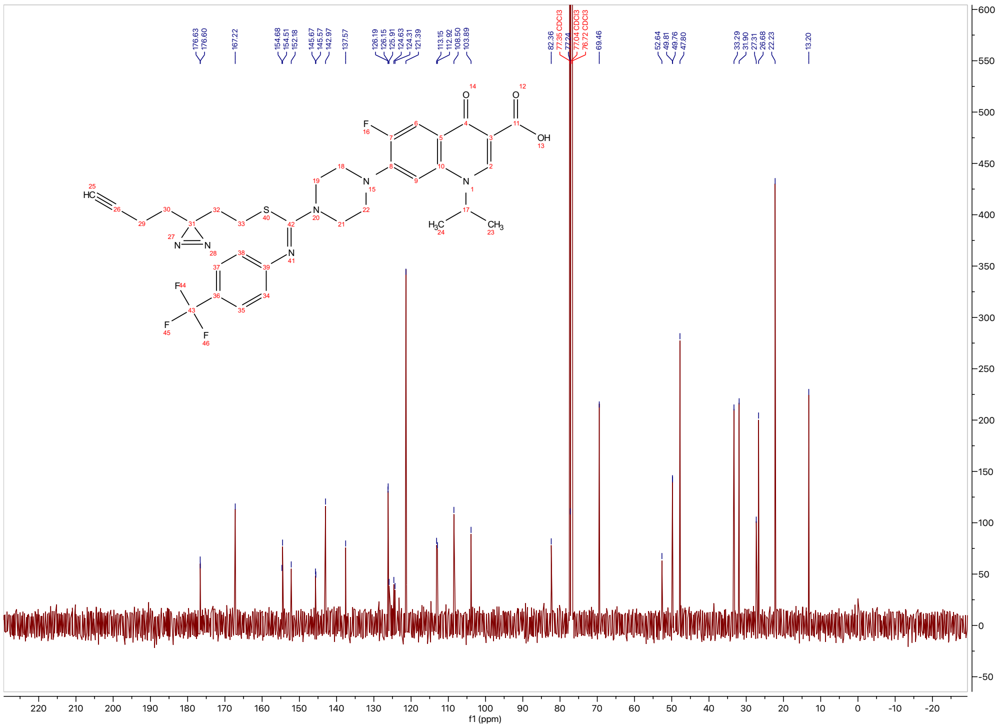

$^{13}C$ NMR (151 MHz, $CDCl_3$) at 298 K of **3**.

**6-fluoro-1-(2-methylbut-3-yn-2-yl)-4-oxo-7-(4-((4-(3-(trifluoromethyl)-3*H*-diazirin-3-yl)benzyl)(4-(trifluoromethyl)phenyl)carbamothioyl)piperazin-1-yl)-1,4-dihydroquinoline-3-carboxylic acid** (**4**)

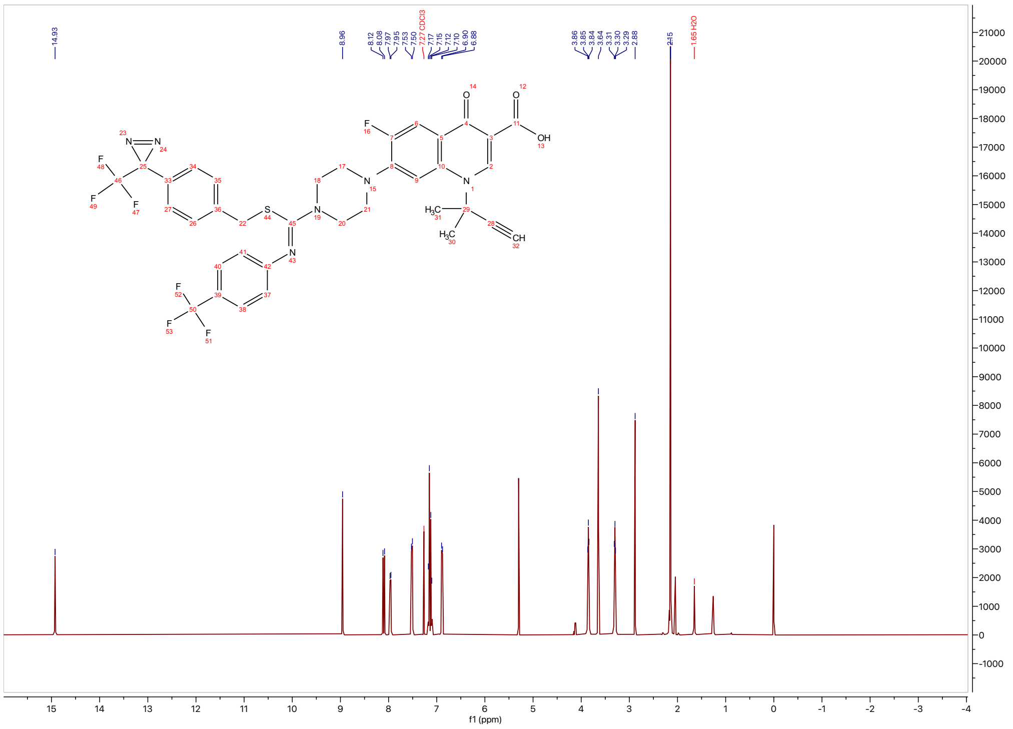

$^{1}H$ NMR (400 MHz, $CDCl_3$) at 298 K of **4**.

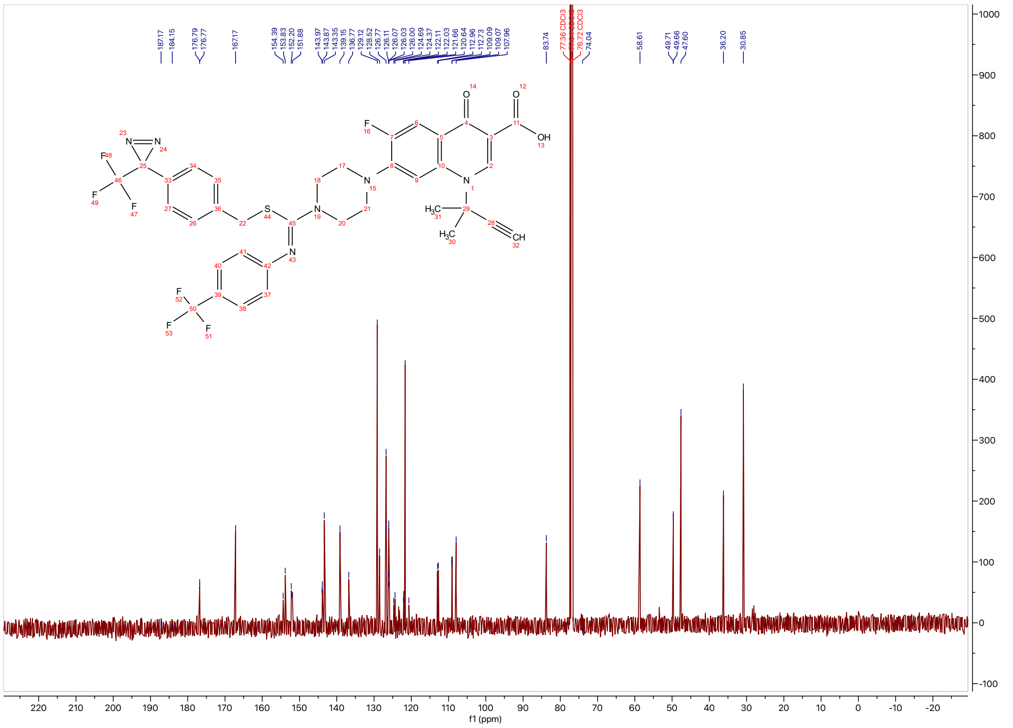

$^{13}C$ NMR (151 MHz, $CDCl_3$) at 298 K of **4**.

**6-Fluoro-4-oxo-7-(4-((prop-2-yn-1-ylthio)((4-(trifluoromethyl)phenyl)imino)methyl)piperazin-1-yl)-1-(4-(3-(trifluoromethyl)-3*H*-diazirin-3-yl)benzyl)-1,4-dihydroquinoline-3-carboxylic acid** (5)

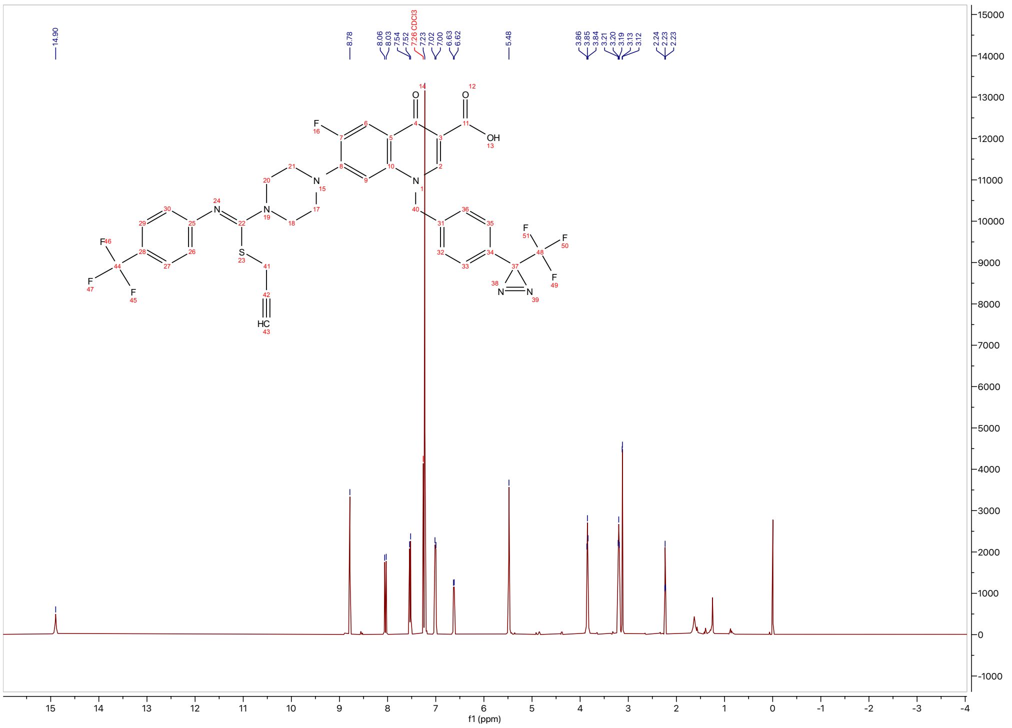

$^{1}$H NMR (400 MHz, $CDCl_3$) at 298 K of **5**.

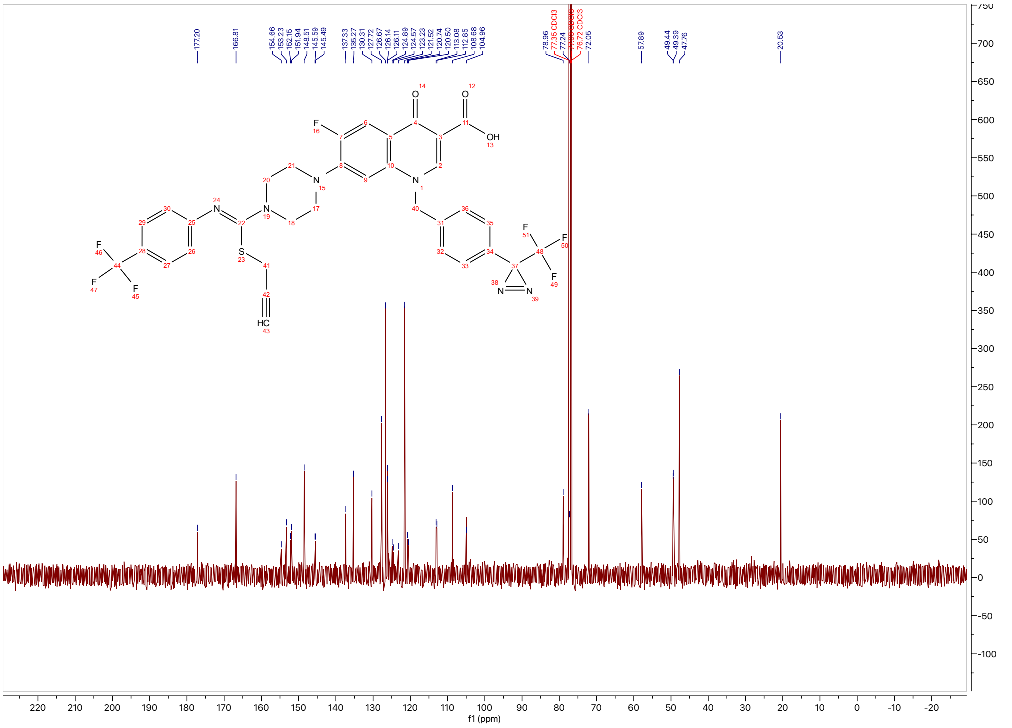

$^{13}$C NMR (151 MHz, $CDCl_3$) at 298 K of **5**.

**1-(2-(3-(but-3-yn-1-yl)-3H-diazirin-3-yl)ethyl)-6-fluoro-4-oxo-7-(4-((4-(trifluoromethyl)phenyl)carbamothioyl)piperazin-1-yl)-1,4-dihydroquinoline-3-carboxylic acid (6)**

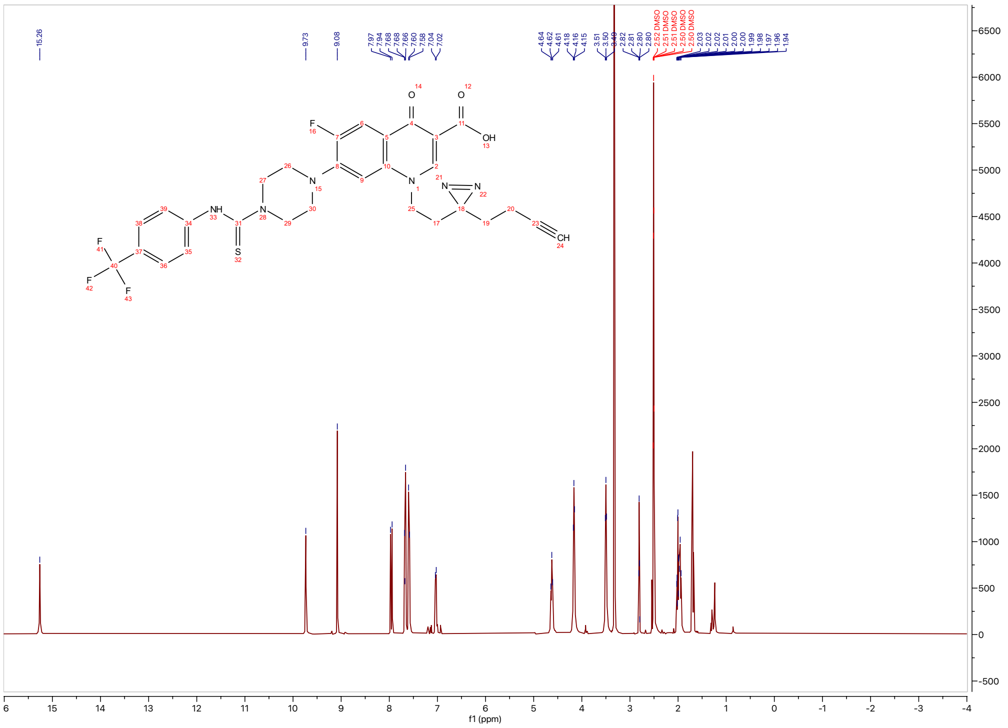

$^1$H NMR (400 MHz, DMSO-$d_6$) at 298 K of **6**.

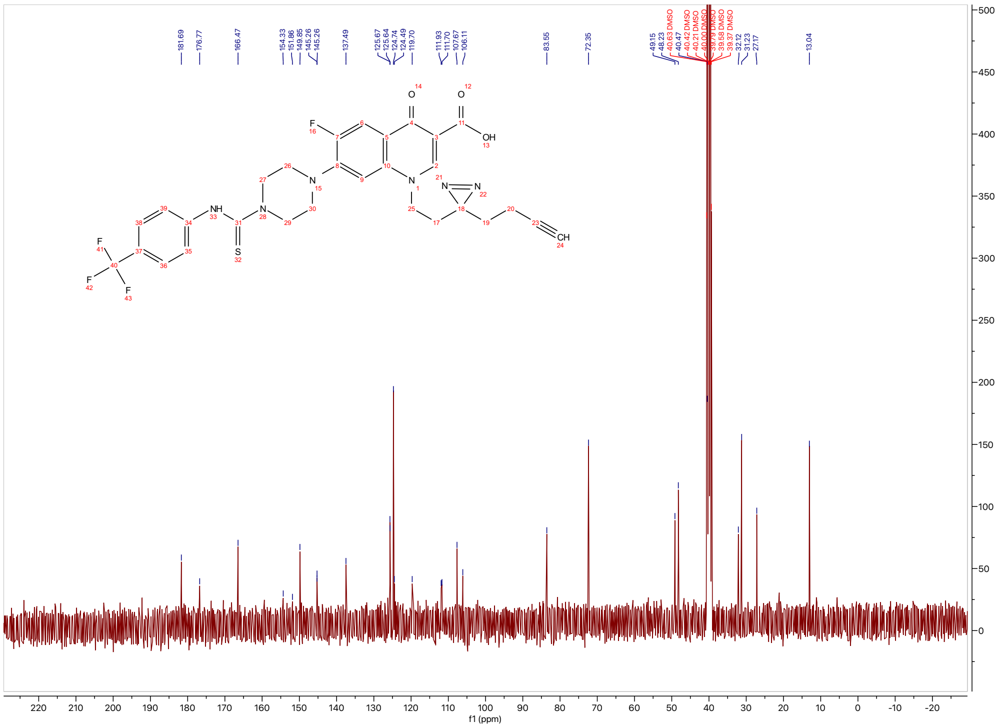

$^{13}$C NMR (151 MHz, DMSO-$d_6$) at 298 K of **6**.

### HPLC Trace of Tested Compounds

#### 7-(4-(((2-(3-(but-3-yn-1-yl)-3H-diazirin-3-yl)ethyl)thio)((4-(trifluoromethyl)phenyl)imino)methyl)piperazin-1-yl)-6-fluoro-1-isopropyl-4-oxo-1,4-dihydroquinoline-3-carboxylic acid (3)

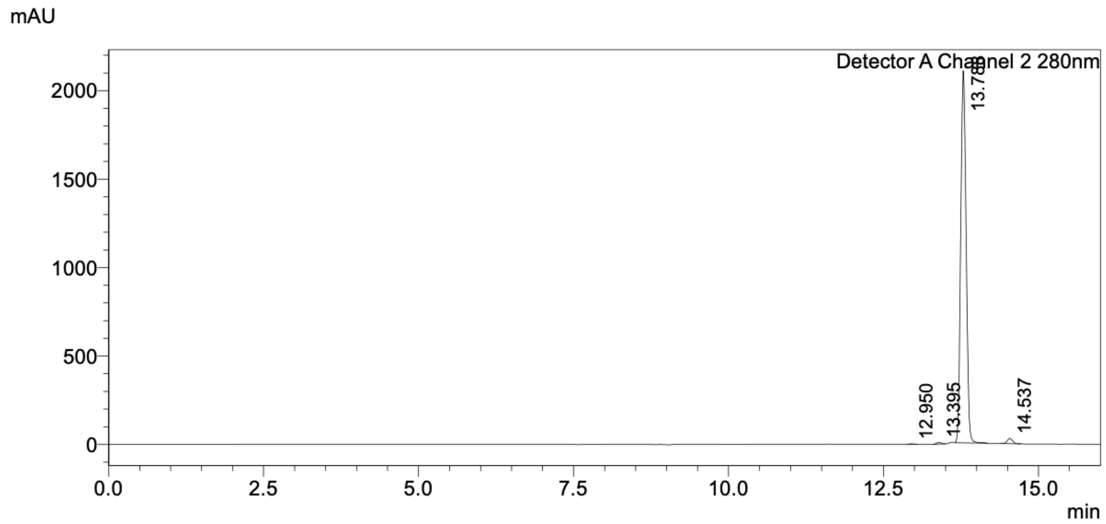

Detector A Channel 2 280nm

| Peak# | Ret. Time | Area | Height | Conc. | Unit | Mark | Name |
|---|---|---|---|---|---|---|---|
| 1 | 12.950 | 13393 | 2309 | 0.103 | | M | |
| 2 | 13.395 | 55715 | 9240 | 0.428 | | M | |
| 3 | 13.788 | 12757319 | 2104517 | 98.087 | | M | |
| 4 | 14.537 | 179741 | 29464 | 1.382 | | M | |
| Total | | 13006167 | 2145531 | | | | |

#### 6-fluoro-1-(2-methylbut-3-yn-2-yl)-4-oxo-7-(4-((4-(3-(trifluoromethyl)-3*H*-diazirin-3-yl)benzyl)(4-(trifluoromethyl)phenyl)carbamothioyl)piperazin-1-yl)-1,4-dihydroquinoline-3-carboxylic acid (4)

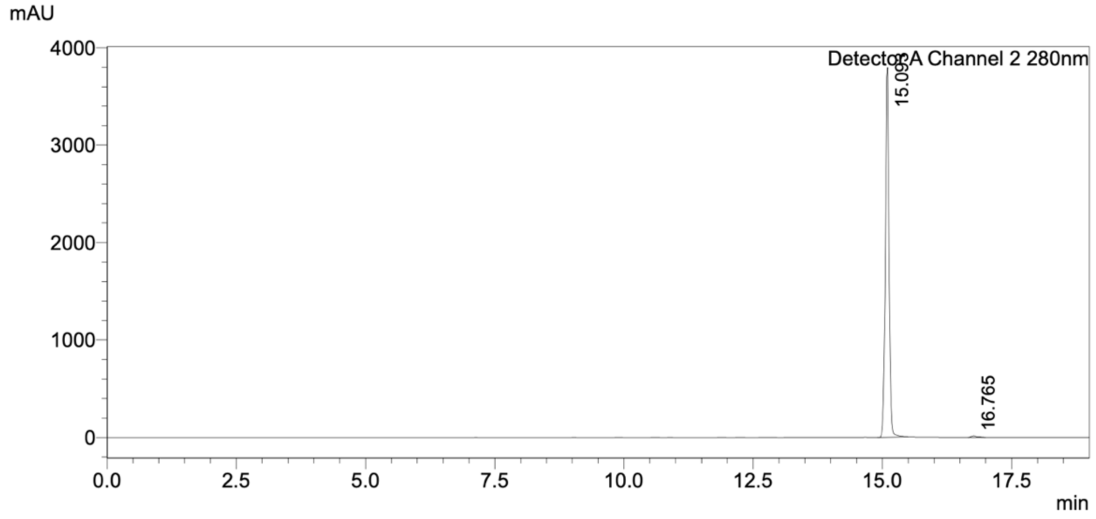

Detector A Channel 2 280nm

| Peak# | Ret. Time | Area | Height | Conc. | Unit | Mark | Name |
|---|---|---|---|---|---|---|---|
| 1 | 15.093 | 18804693 | 3794976 | 99.382 | | M | |
| 2 | 16.765 | 116973 | 15520 | 0.618 | | M | |
| Total | | 18921666 | 3810497 | | | | |

**6-Fluoro-4-oxo-7-(4-((prop-2-yn-1-ylthio)((4-(trifluoromethyl)phenyl)imino)methyl)piperazin-1-yl)-1-(4-(3-(trifluoromethyl)-3*H*-diazirin-3-yl)benzyl)-1,4-dihydroquinoline-3-carboxylic acid** (**5**)

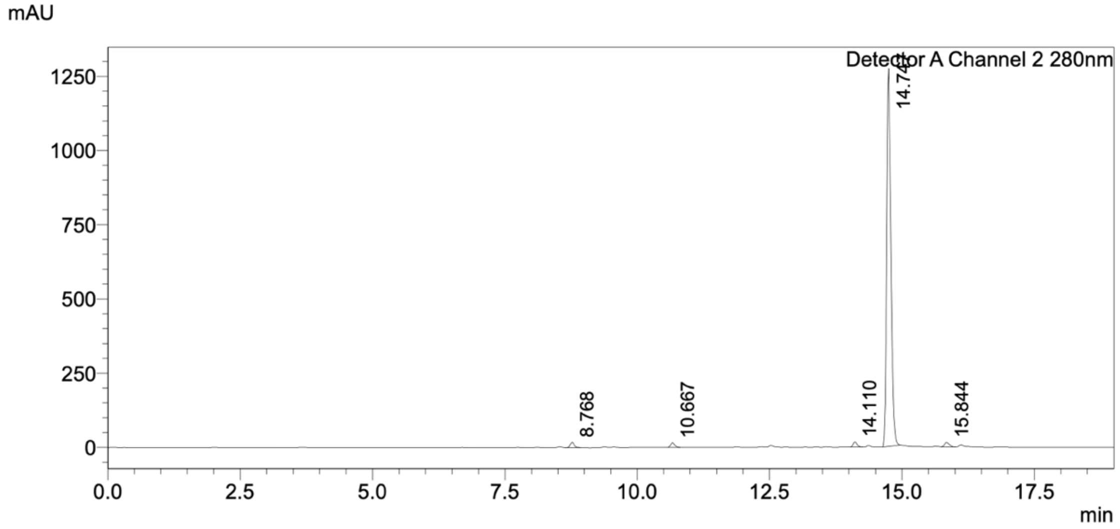

Detector A Channel 2 280nm

| Peak# | Ret. Time | Area | Height | Conc. | Unit | Mark | Name |
|---|---|---|---|---|---|---|---|
| 1 | 8.768 | 86532 | 16937 | 1.162 | | M | |
| 2 | 10.667 | 79052 | 14672 | 1.062 | | M | |
| 3 | 14.110 | 86561 | 16393 | 1.163 | | M | |
| 4 | 14.747 | 7103290 | 1272551 | 95.398 | | M | |
| 5 | 15.844 | 90525 | 14420 | 1.216 | | M | |
| Total | | 7445959 | 1334972 | | | | |

**1-(2-(3-(but-3-yn-1-yl)-3H-diazirin-3-yl)ethyl)-6-fluoro-4-oxo-7-(4-((4-(trifluoromethyl)phenyl)carbamothioyl)piperazin-1-yl)-1,4-dihydroquinoline-3-carboxylic acid** (**6**)

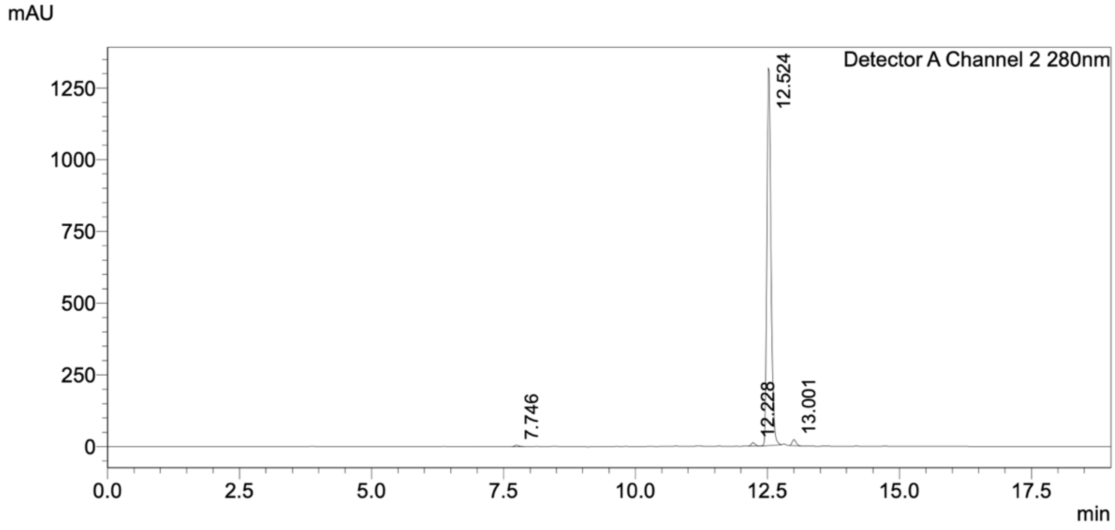

Detector A Channel 2 280nm

| Peak# | Ret. Time | Area | Height | Conc. | Unit | Mark | Name |
|---|---|---|---|---|---|---|---|
| 1 | 7.746 | 29067 | 5503 | 0.392 | | M | |
| 2 | 12.228 | 61232 | 11772 | 0.826 | | M | |
| 3 | 12.524 | 7211131 | 1314410 | 97.311 | | M | |
| 4 | 13.001 | 108993 | 21364 | 1.471 | | M | |
| Total | | 7410422 | 1353049 | | | | |
